# Inhibition of coronaviral exonuclease activity by TRIM-mediated SUMOylation

**DOI:** 10.1101/2025.07.28.667286

**Authors:** Kannan Balakrishnan, Surajit Chakraborty, Cindy Chiang, Caleb Stratton, Anna A. Tumanova, Shaun K. Olsen, Michaela U. Gack

## Abstract

Members of the TRIM E3 ligase family are effectors of the host innate or intrinsic defense against various viruses; however, how specific TRIM proteins antagonize coronavirus infection is still largely elusive. Through an RNAi screen targeting 71 human TRIM genes, we identified multiple TRIM proteins with antiviral or proviral activity against SARS-CoV-2. TRIM32 potently restricted SARS-CoV-2 replication in a RING E3 ligase-dependent but interferon-independent manner. Mechanistically, TRIM32 binds to and SUMOylates the 3′-to-5′ exoribonuclease (ExoN) of NSP14, which is essential for SARS-CoV-2 replication. TRIM32-mediated NSP14 SUMOylation at K9 and K200 inhibits RNA binding and NSP10 cofactor recruitment, respectively, ultimately suppressing ExoN activity. Our study further revealed that NSP14 SUMOylation by TRIM32 and its antiviral activity are broadly conserved for coronaviruses. These results identify the coronaviral NSP14 protein as a direct target of host restriction via SUMOylation, which may uncover novel ways to therapeutically inhibit coronavirus infections in humans.

## INTRODUCTION

The *Coronaviridae* family comprises viruses that are common throughout the world and cause mild to severe respiratory illnesses (*e.g.* bronchitis, pneumonia). Severe acute respiratory syndrome coronavirus 2 (SARS-CoV-2), the causative agent of the COVID-19 pandemic, has caused >7 millions of deaths globally and continues to persist as an endemic pathogen. SARS-CoV-2 is an enveloped, positive-strand RNA virus encoding ∼30 viral proteins – 16 nonstructural (NSP1-NSP16), 4 structural (spike (S), envelope, membrane, and nucleocapsid (N)), and 9 accessory (ORF3a/b, ORF6, ORF7a/b, ORF8, ORF9b/c and ORF10)^1–3^. These viral proteins either have fundamental functions in the viral life cycle (*e.g.* by aiding in virus entry, RNA synthesis, or egress) and/or intricately modulate intrinsic, innate, or adaptive immunity to shape disease outcome. Among the SARS-CoV-2 non-structural proteins is NSP14, which is highly conserved among different coronaviruses, and has two enzymatic activities essential for virus replication: an N-terminal 3′-to-5′ exoribonuclease (ExoN) and a C-terminal N7-methyltransferase (N7-MTase). NSP14-ExoN, together with its cofactor NSP10, catalyzes nucleoside monophosphate excision from RNA, and this activity is crucial for coronaviral proofreading activity and high-fidelity replication^4–9^. Furthermore, the ExoN activity significantly contributes to the resistance of coronaviruses to RNA mutagens (*i.e.* ribonucleoside analogs)^8^. As such, targeting NSP14’s ExoN activity (*e.g.* by blocking the NSP10-NSP14 interaction) is a promising strategy to develop anti-coronaviral therapeutics and sensitizing agents^8^. Whether essential coronaviral enzymatic activities, particularly the ExoN activity of NSP14, are directly antagonized by host-cellular restriction factors is unknown.

*Tripartite motif* (TRIM) proteins are a family of E3 ligases (>70 members encoded in the human genome) that play crucial roles in host restriction of virus infections^10–12^. In turn, numerous RNA viruses and DNA viruses (*e.g.* influenza, flaviviruses, herpesviruses) antagonize distinct TRIM proteins to effectively replicate and disseminate in the host^11,13^. Although new mechanisms for TRIM proteins are still emerging, TRIM proteins have been shown to act antiviral via three principal modes of action: *1)* potentiation of type I and III interferon (IFN) responses; *2)* modulation of autophagic or apoptotic host processes; and *3)* direct antagonism of viral protein functions or virus gene transcription, impeding key steps in the viral replication cycle. Notably, several TRIM proteins restrict viruses via more than one of these mechanisms. For instance, TRIM25 and TRIM65 potentiate the activation of cytoplasmic innate sensing pathways (*i.e.* RIG-I-like receptor signaling^14,15^), but also directly impede viral factors or modulate autophagy in certain contexts^12,16,17^. Similarly, TRIM5α blocks HIV-1 via an interaction with the viral capsid, but also boosts intrinsic immunity via catalysis of non-degradative K63-linked ubiquitination^18^. Another emerging key principle is that specific TRIM proteins can restrict different viruses via unique mechanisms and distinct protein-protein interactions^12^. Compared to our knowledge about antiviral roles for TRIM proteins, significantly less is known about their proviral functions. Moreover, little is still known about how TRIM proteins impact the replication of coronaviruses, in particular SARS-CoV-2.

In this study, we identified through an unbiased RNAi screen multiple TRIM proteins with potent anti-or pro-viral activities against SARS-CoV-2 in human cells. Our molecular characterization studies on TRIM32 revealed a unique mechanism of virus restriction whereby the coronaviral NSP14-ExoN is a direct target of TRIM32-mediated SUMOylation, blocking viral ExoN activity.

## RESULTS

### RNAi screen identifies antiviral and proviral TRIM proteins in SARS-CoV-2 infection

To identify TRIM proteins that modulate SARS-CoV-2 replication, we performed an RNAi screen targeting 71 human TRIM genes to evaluate the effect of TRIM knockdown on virus replication (**Figures 1A and 1B**). Individual TRIM genes were silenced by transient transfection in Caco-2 (human intestinal epithelial) cells engineered to stably express human ACE2 (Caco-2-hACE2) followed by infection with recombinant SARS-CoV-2 expressing green fluorescent protein (EGFP-rSARS-CoV-2). The real-time replication kinetics of EGFP-rSARS-CoV-2 was assessed every hour over a 72-h period by monitoring EGFP expression in live cells. Scrambled non-targeting siRNA (si.C) served as control. Furthermore, siRNA targeting LY6E (lymphocyte antigen 6 family member E), which inhibits SARS-CoV-2 membrane fusion^19,20^, and siRNA targeting TMPRSS2 (transmembrane protease serine 2), a key enzyme for efficient SARS-CoV-2 infection^21^, were included for comparison. Silencing of seven TRIM proteins (*i.e.* TRIM6, TRIM18, TRIM21, TRIM32, TRIM41, TRIM65 and TRIM70) showed robustly enhanced EGFP-rSARS-CoV-2 replication as compared to si.C transfection, even more strongly compared to LY6E knockdown, suggesting that these TRIM proteins restrict SARS-CoV-2 replication (**Figures 1B and 1C**). Conversely, cells individually depleted of TRIM4, TRIM5, TRIM40, TRIM44, TRIM45, TRIM55, and TRIM59 showed a decrease in EGFP-rSARS-CoV-2 replication compared to si.C transfection, and the effect was similar to that of TMPRSS2 silencing, suggesting proviral roles of these TRIM proteins in SARS-CoV-2 infection (**Figures 1B and 1D**). Of note, knockdown of individual TRIM proteins had little to no cytotoxic effects (**Figure S1A**).

**Figure 1.**
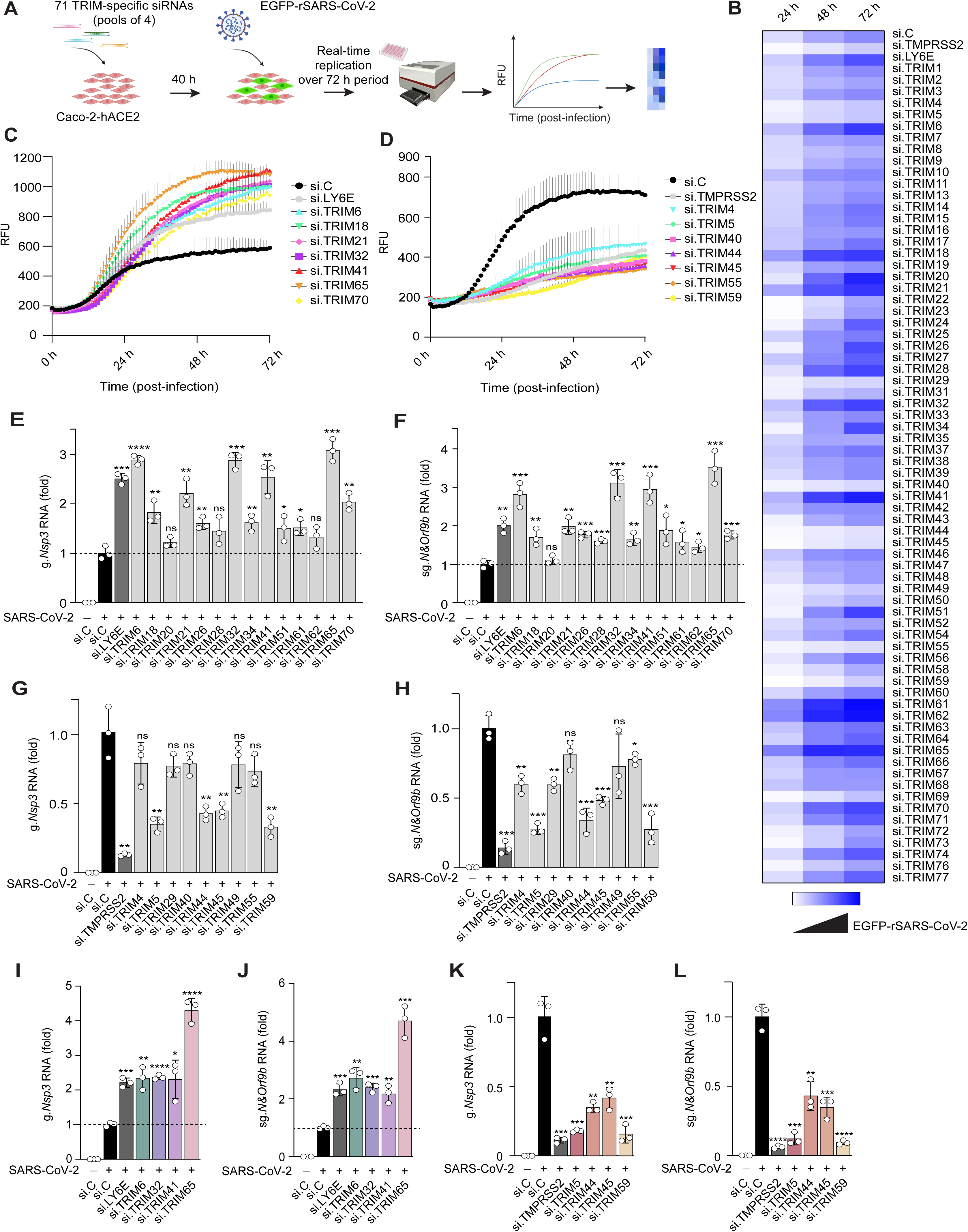
Antiviral and proviral TRIM proteins in SARS-CoV-2 infection. (**A**) Schematic representation of the RNAi screen targeting 71 human TRIM genes to test the effect of TRIM gene silencing on SARS-CoV-2 replication. Caco-2 cells stably expressing human ACE2 (Caco-2-hACE2) were transfected for 40 h with the indicated TRIM-specific siRNAs (pool of 4 different siRNAs per TRIM), or non-targeting control siRNA (si.C), and then either mock-treated (−) or infected with EGFP-rSARS-CoV-2 (strain K49; MOI 0.02). siRNAs targeting TMPRSS2 (proviral) and LY6E (antiviral) were included as controls. Virus replication kinetics in live cells was assessed every hour for a 72-h period by measuring relative fluorescence units (RFU) for EGFP. Parts of this figure were created using Biorender.com. (**B**) Heat map representation of the EGFP-rSARS-CoV-2 replication kinetics from the TRIM RNAi screen in (A). (**C** and **D**) EGFP-rSARS-CoV-2 replication kinetics for selected TRIM proteins identified in the RNAi screen showing antiviral (C) or proviral (D) activities, presented as RFU. Values were auto-scaled to the average values for mock-infected cells at 0 h, set to 200. (**E—H**) SARS-CoV-2 genomic RNA (gRNA; *Nsp3*) (E and G) and subgenomic RNA (sgRNA; *N&Orf9b*) (F and H) in Caco-2-hACE2 cells that were transfected for 40 h with either si.C or the indicated TRIM-specific siRNAs and then either mock-treated (−) or infected with SARS-CoV-2 (WA1, MOI 0.02) for 48 h, determined by RT-qPCR. si.LY6E and si.TMPRSS2 were included as controls. (**I—L**) SARS-CoV-2 genomic RNA (g.*Nsp3*) (I and K) and subgenomic RNA (sg.*N&Orf9b*) (J and L) in Calu-3 cells that were transfected for 40 h with either si.C or siRNAs targeting the indicated human TRIM proteins or controls, and then either mock-treated (−) or infected with SARS-CoV-2 (WA1, MOI 0.02) for 48 h, determined by RT-qPCR. Data are representative of one RNAi screen (conducted in triplicates (B—D)) or at least two independent experiments (mean ± s.d. of n = 3 biological replicates (E—L)). *p < 0.05, **p < 0.01, ***p < 0.001, ****p < 0.0001 (Student’s *t* test). ns, not significant. See also Figure S1.

To validate the top antiviral and proviral TRIM proteins that emerged from our screen, we tested the effect of their depletion on SARS-CoV-2 subgenomic and genomic RNA (sgRNA and gRNA) production in Caco-2-hACE2 cells. Several other TRIM genes were silenced in these experiments for comparison (**Figures 1E—H and Figure S1B**). Knockdown of endogenous TRIM6, TRIM32, TRIM41 and TRIM65 enhanced SARS-CoV-2 gRNA and sgRNA expression to comparable or even stronger levels than did LY6E silencing (**Figures 1E and 1F**). Depletion of endogenous TRIM5, TRIM44, TRIM45 and TRIM59 strongly reduced SARS-CoV-2 gRNA and sgRNA levels (as compared to si.C transfection), however not as efficiently as did TMPRSS2 knockdown (**Figures 1G and 1H**). Of note, silencing of the other TRIM proteins led to only minor or no significant changes in viral RNA levels as compared to si.C transfection (**Figures 1E—H**), hinting at the possibility that these TRIMs may regulate SARS-CoV-2 replication at a step post-RNA synthesis, which warrants further investigation. Individual depletion of endogenous TRIM6, TRIM32, TRIM41 and TRIM65 in Calu-3 (human airway epithelial) cells also led to significantly enhanced SARS-CoV-2 RNA levels (**Figures 1I and 1J and Figure S1C**), while knockdown of TRIM5, TRIM44, TRIM45 and TRIM59 strongly reduced SARS-CoV-2 RNA expression (**Figures 1K and 1L and Figure S1C**). Of note, silencing of some TRIM proteins, in particular TRIM65, an activator of type I IFN responses^15,22^, influenced SARS-CoV-2 replication in Caco-2 versus Calu-3 cells to a different extent, suggesting cell type-specific differences in the potency of IFN regulation by TRIM65. Altogether, these data identify several TRIM proteins as anti-or pro-viral factors modulating SARS-CoV-2 replication.

### TRIM32 restricts SARS-CoV-2 in an E3 ligase-dependent but IFN-independent manner

We focused our functional characterization studies on TRIM32 which, when silenced, showed a strong enhancement of SARS-CoV-2 replication in different cell types, and whose antiviral function against coronaviruses is unknown. Utilizing CRISPR-Cas9 technology, we generated *TRIM32*-knockout (KO) Caco-2 single-cell clones and tested SARS-CoV-2 replication in these cells compared to wild-type (WT) control cells (**Figures 2A and 2B**). Consistent with our data using TRIM32-specific siRNAs, *TRIM32*-KO cells showed markedly increased viral replication over a period of 72 h as compared to WT control cells (**Figure 2C**). In accord, SARS-CoV-2 gRNA/sgRNA and viral protein (*i.e.* S and N) expression were strongly enhanced in *TRIM32*-KO cells as compared to infected WT cells (**Figures 2D—F**).

**Figure 2.**
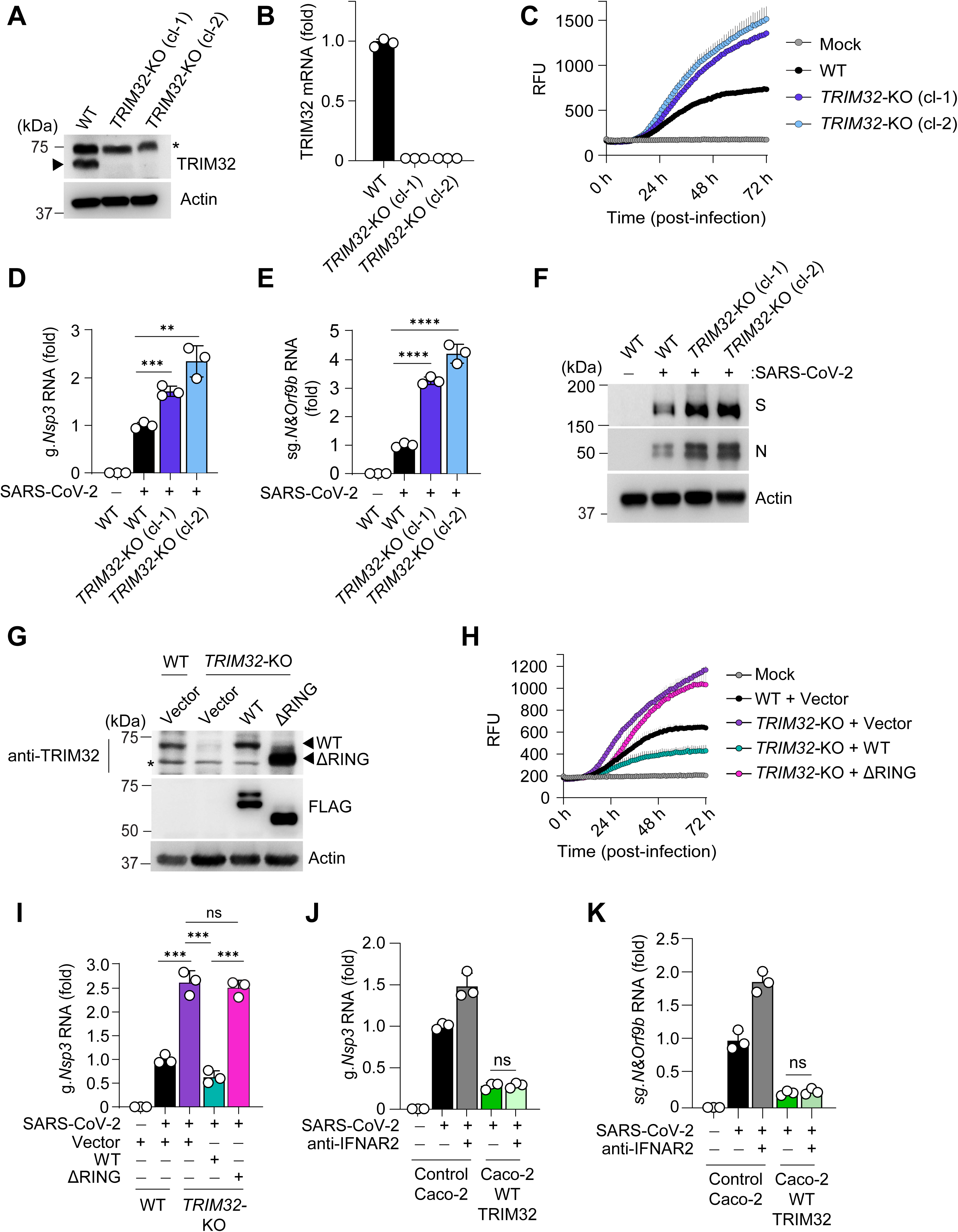
**TRIM32 restricts SARS-CoV-2 replication in a RING-dependent but IFN-independent manner** (**A**) Validation of CRISPR *TRIM32-*KO Caco-2 cells. TRIM32 protein abundance in *TRIM32*-KO Caco-2 single cell clones (cl-1 and cl-2) and WT control cells, determined in the whole cell lysates (WCLs) by immunoblot (IB) analysis with anti-TRIM32 and anti-Actin (loading control). Asterisk indicates nonspecific band. (**B**) Validation of CRISPR *TRIM32-*KO Caco-2 cells (cl-1 and cl-2) by assessing *TRIM32* transcripts by RT-qPCR. Values indicate relative mRNA (fold), normalized to cellular *HPRT1*. (**C**) Virus replication kinetics over a 72-h period in *TRIM32*-KO and WT Caco-2 cells that were either mock-treated or infected with EGFP-rSARS-CoV-2 (strain K49; MOI 0.02), determined by measuring RFU for EGFP. Values were auto-scaled to the average values for mock-infected cells at 0 h, set to 200. (**D** and **E**) RT-qPCR analysis of SARS-CoV-2 g.*Nsp3* (D) and sg.*N&Orf9b* (E) transcripts in *TRIM32*-KO and WT Caco-2 cells that were either mock-treated (−) or infected with SARS-CoV-2 (strain WA1; MOI 0.02) for 48 h. (**F**) SARS-CoV-2 spike (S) and nucleocapsid (N) protein abundances in *TRIM32*-KO and WT Caco-2 cells that were mock-treated (−) or infected as in (D), determined in the WCLs by IB with anti-S and anti-N. (**G**) TRIM32 protein expression in stable reconstituted *TRIM32*-KO Caco-2 cells that inducibly (via doxycycline (DOX; 2 ug/mL) treatment for 24 h) expressed either empty vector or FLAG-tagged TRIM32 WT or ΔRING, determined in the WCLs by IB with anti-TRIM32, anti-FLAG, and anti-Actin (loading control). WT Caco-2 cells expressing empty vector served as control. Asterisk indicates nonspecific band. (**H**) Virus replication kinetics over a 72-h period in *TRIM32*-KO Caco-2 cells that were reconstituted as in (G) and then either mock-treated or infected with EGFP-rSARS-CoV-2 (strain K49; MOI 0.02), determined by measuring RFU for EGFP. Values were auto-scaled as in (C). WT Caco-2 cells expressing empty vector served as control. (**I**) SARS-CoV-2 g.*Nsp3* transcripts in *TRIM32*-KO Caco-2 cells reconstituted as indicated and either mock-treated (−) or infected with SARS-CoV-2 (strain WA1; MOI 0.02) for 48 h, determined by RT-qPCR. WT Caco-2 cells expressing empty vector served as control. (**J** and **K**) SARS-CoV-2 g.*Nsp3* and sg.*N&orf9b* transcripts in stable Caco-2 cells that inducibly (via DOX (2 ug/mL) treatment) expressed empty vector (control) or TRIM32 WT and were infected with SARS-CoV-2 (strain WA1; MOI 0.05) for 48 h in the absence or presence of anti-IFNAR2 antibody (2 µg/mL), determined by RT-qPCR. Data are representative of at least two independent experiments (mean ± s.d. of n = 3 biological replicates (B—E, H—K)). **p < 0.01, ***p < 0.001, ****p < 0.0001 (Student’s *t* test). ns, not significant. See also Figure S2.

TRIM proteins are characterized by an E3 ligase activity conferred by the N-terminal RING domain which, for many TRIM proteins, is crucial for antiviral activity^10,11^. To determine if the E3 ligase activity of TRIM32 is important for SARS-CoV-2 restriction, we generated stable Caco-2 cells inducibly expressing either empty vector, TRIM32 WT or the ΔRING mutant (**Figure S2A**). Cells expressing TRIM32 WT, but not cells expressing TRIM32ΔRING, effectively suppressed SARS-CoV-2 replication (**Figures S2B—D**). In accord, stable expression of TRIM32 WT, but not ΔRING, suppressed viral S and N protein expression (**Figure S2E**). These results were corroborated in *TRIM32*-KO cells stably reconstituted with either TRIM32 WT or ΔRING, as cells complemented with TRIM32 WT, but not ΔRING, effectively restricted SARS-CoV-2 replication and viral RNA production (**Figures 2G—I**). These results indicate that TRIM32-mediated suppression of SARS-CoV-2 replication requires the RING E3 ligase activity.

As TRIM32 has been reported to regulate type I IFN induction and thereby antiviral defenses (of note, however, not in the context of coronavirus infection)^23^, we next determined whether TRIM32 impedes SARS-CoV-2 replication in an IFN-dependent or -independent manner. We used a blocking antibody targeting human IFNAR2 (IFN-α/β receptor subunit 2) to ablate the effects of type I IFN signaling. Effective IFNAR-signal blockage by anti-IFNAR2 was confirmed by measuring virus-induced ISG (*i.e.* IFIT2 and ISG15) mRNA expression (**Figures S2F and S2G**). We found that stable TRIM32 expression potently suppressed SARS-CoV-2 RNA production in the presence or absence of anti-IFNAR2 (**Figures 2J and 2K**). In accord, immunofluorescence (IF) microscopy analysis showed that EGFP-rSARS-CoV-2 replication was equally diminished by stable TRIM32 expression in Caco-2 cells with or without anti-IFNAR2 administration (**Figure S2H**). Collectively, these data indicate that TRIM32 inhibits SARS-CoV-2 replication in an IFN-independent but RING E3 ligase-dependent fashion.

### TRIM32 interacts with NSP14 and catalyzes its SUMOylation

We postulated that TRIM32 may directly antagonize the function of specific SARS-CoV-2 protein(s) that are essential for virus replication. We first examined whether endogenous TRIM32 co-localizes with the coronaviral replication-transcription complex (RTC), an assembly of NSPs required for coordinating efficient viral genome replication and RNA transcription. Laser scanning confocal microscopy analysis showed that whereas endogenous TRIM32 was localized throughout the cytoplasm in uninfected cells (of note, a minor portion of TRIM32 localized to the nucleus as previously reported^24^), it showed a prominent co-localization with viral dsRNA, indicative of viral RTCs, in SARS-CoV-2-infected cells (**Figures S3A and S3B**).

Published proteomics screens suggested that TRIM32 interacts with SARS-CoV-2 NSP14 and NSP16^25,26^; however, whether these interactions occur in physiological contexts and are relevant for modulating virus replication, is unknown. NSP14 harbors two essential enzymatic activities (*i.e.* ExoN and N7-MTase) and is essential for SARS-CoV-2 replication^7,8^, while NSP16 of coronaviruses is critical for immune escape of IFN responses^27–32^. Co-immunoprecipitation (Co-IP) assay showed efficient binding of V5-tagged TRIM32 to FLAG-tagged NSP14 and NSP16. In contrast, TRIM32 did not interact with NSP2, NSP10, or NSP15, which were included as controls, under the same conditions (**Figure S3C**).

Since our data indicated that TRIM32 inhibits SARS-CoV-2 replication in an IFN-independent manner, and because NSP16 functions primarily in innate immune escape, we focused on examining TRIM32’s role in targeting NSP14, which is a key enzyme essential for SARS-CoV-2 replication. Consistent with our binding experiments assessing ectopic proteins, we found that endogenous TRIM32 readily interacted with NSP14 during authentic SARS-CoV-2 infection (**Figure 3A**).

**Figure 3.**
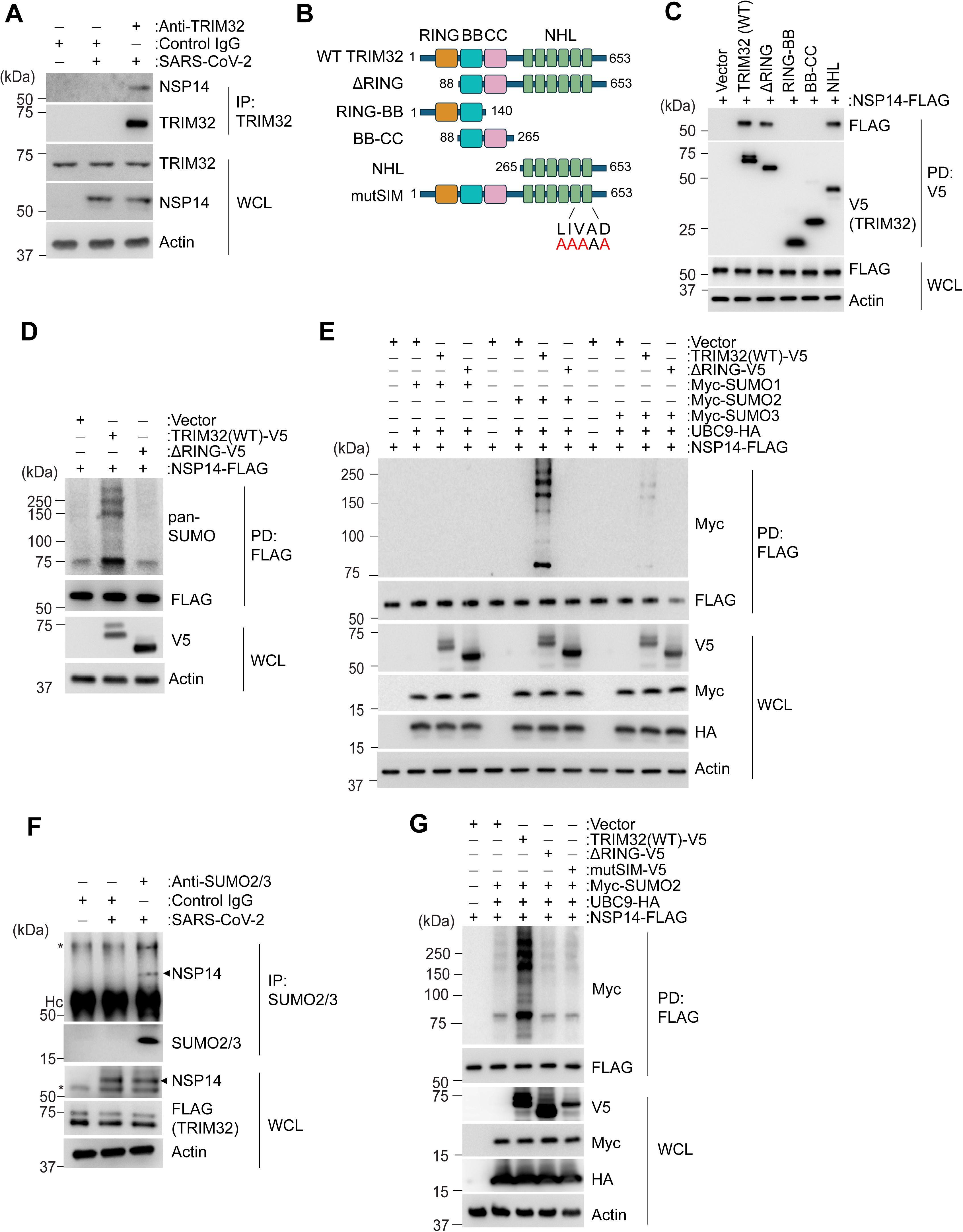
**TRIM32 binds to and SUMOylates SARS-CoV-2 NSP14** (**A**) Endogenous TRIM32 binding to NSP14 in Caco-2 cells that were either mock-treated (−) or infected with SARS-CoV-2 (strain WA1; MOI 1) for 48 h, determined by IP with anti-TRIM32 and IB with anti-NSP14. (**B**) Schematic representation of TRIM32’s domain organization as well as the truncation constructs used in the interaction-mapping experiments. RING, really interesting new gene; BB, B-box; CC, coiled-coil; NHL, NCL-1, HT2A, Lin-41 domain; mutSIM, a mutant of TRIM32 in which the SIM motif located in the NHL domain is mutated. Mutated residues are shown in red. Numbers indicate amino acids. (**C**) Binding of FLAG-NSP14 (SARS-CoV-2) to the indicated V5-tagged TRIM32 truncation proteins in transiently transfected HEK293T cells, assessed by PD with anti-V5 (PD: V5), followed by IB with anti-FLAG. WCLs were probed with anti-FLAG and anti-Actin (loading control). (**D**) SUMOylation of FLAG-tagged NSP14 (SARS-CoV-2) in HEK293T cells that were co-transfected for 40 h with V5-tagged TRIM32 WT or ΔRING, determined by PD: FLAG and IB with anti-pan-SUMO. WCLs were probed by IB with anti-V5 and anti-Actin (loading control). (**E**) SUMOylation of FLAG-tagged NSP14 (SARS-CoV-2) in transiently transfected HEK293T cells that co-expressed either empty vector or V5-tagged TRIM32 WT or ΔRING, along with UBC9-HA and Myc-tagged SUMO1, SUMO2 or SUMO3, determined by PD: FLAG and IB with anti-Myc and anti-FLAG. Expression of co-transfected proteins was determined in the WCLs using the indicated antibodies. (**F**) SUMOylation of NSP14 in stable Caco-2 cells that were infected for 32 h with SARS-CoV-2 (strain WA1; MOI 2) and then treated for 12 h with DOX (2 µg/mL) to inducibly express FLAG-tagged TRIM32, assessed by denaturing IP with anti-SUMO2/3 and IB with anti-NSP14. WCLs were probed by IB with the indicated antibodies. Hc, antibody heavy chain. Asterisk indicates nonspecific band. (**G**) SUMOylation of FLAG-tagged NSP14 (SARS-CoV-2) in HEK293T cells that were transfected for 40 h with either empty vector or V5-tagged TRIM32 WT or mutants (ΔRING or mutSIM) together with Myc-tagged SUMO2 and HA-tagged UBC9, determined by PD: FLAG and IB with anti-Myc. WCLs were probed by IB with the indicated antibodies. Data (A and C—G) are representative of at least two independent experiments. See also Figure S3.

Human TRIM32 is a 653-amino-acid long protein that harbors, besides an N-terminal RING domain, B-box (BB) and coil-coiled (CC) domains followed by an NHL repeat domain at the C-terminal end (**Figure 3B and Figure S3D**). Biochemical domain-mapping studies showed that NSP14 bound to the C-terminal NHL domain of TRIM32 as efficiently as it interacted with TRIM32 WT or the ΔRING mutant. By contrast, NSP14 did not bind to the RING-BB and BB-CC of TRIM32 **(Figure 3C**). This indicated that the NHL domain of TRIM32 mediates its binding to NSP14.

Our data showed that NSP14’s protein abundance was unaffected by the presence of ectopic TRIM32 expression (**Figure 3C**), suggesting that TRIM32 does not induce NSP14 degradation. Consistent with this concept, ectopic expression of TRIM32 did not potentiate NSP14 ubiquitination (**Figure S3E**), strengthening the proposal that TRIM32 does not trigger ubiquitination-dependent degradation of NSP14. Interestingly, we found that ectopic expression of TRIM32, but not of the ΔRING mutant, robustly induced SUMOylation of affinity-purified NSP14 when probed with a pan-SUMO antibody (*i.e.* an antibody recognizing SUMO1, 2 and 3), where NSP14 showed a high-molecular-weight SUMO modification indicative of poly-SUMO chains (**Figure 3D**). Co-expression of SUMOylation machinery components, specifically UBC9 (also known as UBE2I; the sole E2 enzyme in the SUMOylating pathway) and SUMO2, induced robust NSP14 SUMOylation by TRIM32. By contrast, no NSP14 SUMOylation was observed with the TRIM32 ΔRING mutant, or when SUMO1 or SUMO3 was co-expressed with WT TRIM32 (**Figure 3E**). Importantly, we also readily detected NSP14 SUMOylation during authentic SARS-CoV-2 infection (**Figure 3F**).

To further corroborate whether TRIM32 functions as a *bona fide* SUMO E3 ligase, we tested whether TRIM32 interacts with the E2 enzyme UBC9. Co-IP assay showed that UBC9 efficiently bound to TRIM32 WT and the RING-BB, but not to TRIM32ΔRING or the BB-CC and NHL truncation proteins, indicating that UBC9 is recruited to the RING domain of TRIM32 (**Figure S3F**). Another hallmark of SUMO E3 ligases is the presence of one or more SUMO-interacting motifs (SIMs), which mediate non-covalent binding of SUMO proteins to enhance E3 SUMOylation potency. In most cases, this SIM-mediated interaction helps position the SUMO-loaded E2 enzyme for efficient transfer of SUMO to the substrate^33^. Sequence prediction analysis suggested a putative SIM motif (^589^LIVAD^593^) in the NHL domain of TRIM32 (**Figure 3B**). Functional assessment of NSP14 SUMOylation showed that, unlike WT TRIM32, a mutant in which the SIM motif is mutated (^589^LIVAD^593^ è ^589^AAAAA^593^) (termed ‘mutSIM TRIM32’), did not induce NSP14 SUMOylation (**Figures 3B and 3G**). Altogether, these results indicate that TRIM32, previously known to only mediate ubiquitination events, is a SUMO E3 ligase that catalyzes poly-SUMO2-modifications on NSP14.

### NSP14 is SUMOylated at K9 and K200 in the ExoN domain

To identify the specific residues in NSP14 that undergo SUMOylation by TRIM32, we performed liquid chromatography tandem mass spectrometry (LC-MS/MS) analysis of affinity-purified SARS-CoV-2 NSP14 in cells co-expressing TRIM32, UBC9, and the SUMO2-T91R mutant which allows for capturing the K-G-G signature of SUMOylated substrates^34,35^ (**Figure 4A**). This identified two SUMOylation sites, K9 and K200, located in the N-terminal ExoN domain of NSP14 (**Figure 4B**). Biochemical characterization of NSP14 mutant proteins in which the SUMOylation sites were mutated —either individually (K9R or K200R) or combined (K9R/K200R)— showed that single mutations reduced NSP14 SUMOylation, while the combined mutation of K9 and K200 near-abolished NSP14 SUMOylation (**Figure 4C**). Importantly, the NSP14 K9R, K200R and K9R/K200R mutants bound to TRIM32 as efficiently as WT NSP14 (**Figure 4D**). Taken together, these data identify the ExoN domain as a target of TRIM32-mediated SUMOylation where K9 and K200 serve as the primary SUMOylation sites.

**Figure 4.**
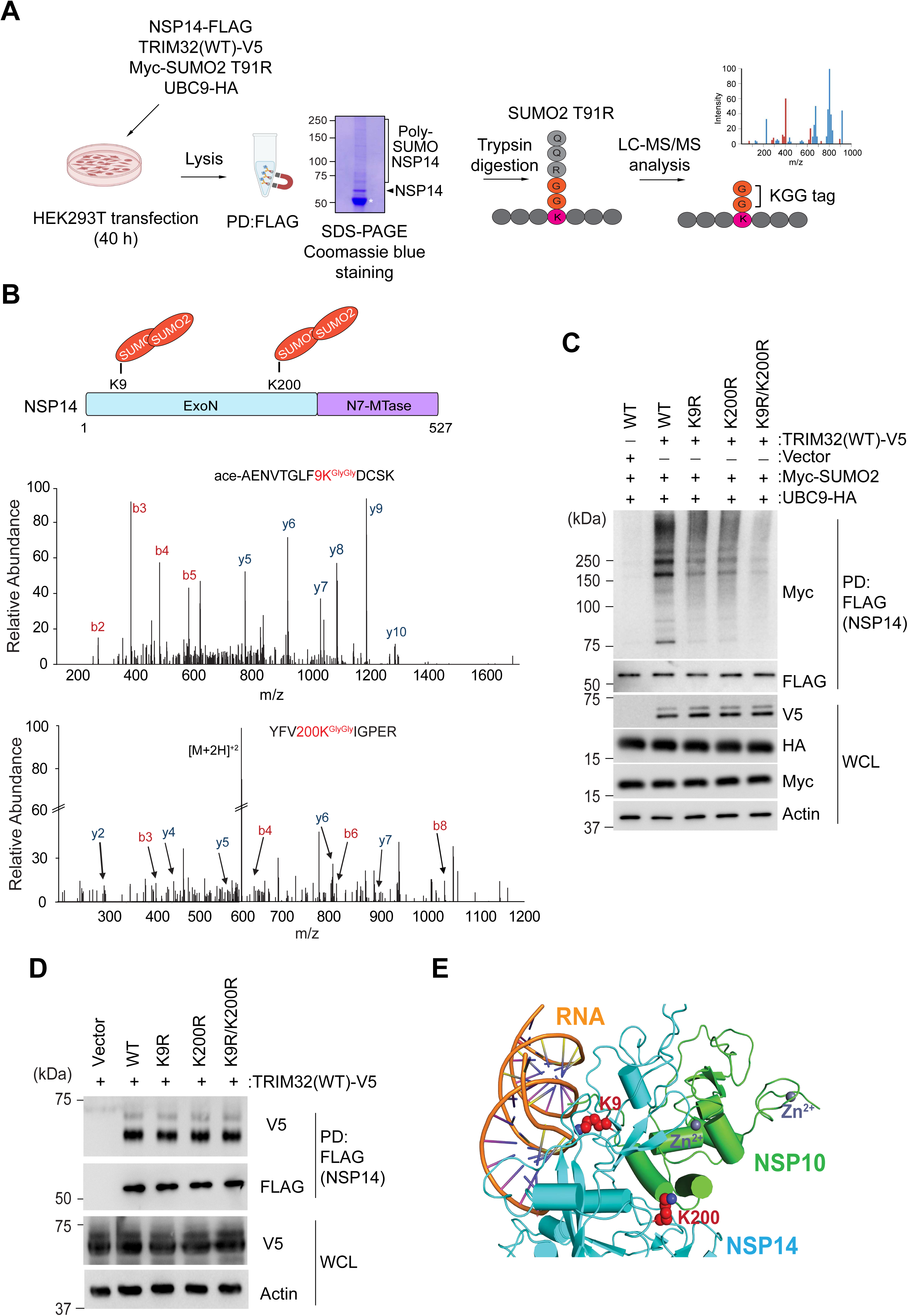
**The NSP14 ExoN domain is SUMOylated at K9 and K200** (**A**) Schematic representation of the approach to identify the SUMOylation sites in NSP14 of SARS-CoV-2 by mass spectrometry (MS) analysis. HEK293T cells were transiently transfected with FLAG-tagged NSP14, V5-tagged TRIM32, HA-tagged UBC9, and Myc-tagged SUMO2 T91R mutant. 40 h later, NSP14 was affinity-purified by PD: FLAG, followed by SDS-PAGE and Coomassie blue staining. Unmodified NSP14 (arrowhead) and modified NSP14 versions are indicated (center). NSP14 protein was subjected to digestion using trypsin, and peptides modified by SUMO2 T91R Lys-Gly-Gly (KGG) signatures were analyzed using LC-MS/MS. Asterisk indicates antibody heavy chain. Parts of this figure were created using Biorender.com. (**B**) Top: Schematic representation of SARS-CoV-2 NSP14 domain organization with the SUMOylated lysines (Ks) identified by MS analysis indicated. ExoN, exonuclease domain; N7-MTase, N7-methyltransferase. Bottom: Mass spectra of the tryptic peptides of affinity-purified FLAG-tagged NSP14 (SARS-CoV-2) that identified SUMOylation at K9 and K200, with designated b-and y-ions. Ace denotes acetylated N-terminus. (**C**) SUMOylation of FLAG-tagged NSP14 (SARS-CoV-2) WT or indicated mutants in transiently transfected HEK293T cells that were co-transfected for 40 h with either empty vector or V5-tagged TRIM32 together with HA-tagged UBC9 and Myc-tagged SUMO2, determined by PD: FLAG and IB with anti-Myc. WCLs were probed by IB with the indicated antibodies. (**D**) Binding of FLAG-tagged NSP14 (SARS-CoV-2) WT or mutants in transiently transfected HEK293T cells that were co-transfected for 40 h with V5-tagged TRIM32, determined by PD: FLAG and IB with anti-V5. WCLs were probed by IB with the indicated antibodies. (**E**) Representation of the cryo-EM structure of the SARS-CoV-2 NSP10-NSP14-RNA complex (PDB: 7N0B) with NSP14 (cyan), RNA (orange), and NSP10 (green). K9 and K200 are depicted in red using spheres. Bound zinc ions are depicted as grey spheres. Data are representative of at least two independent experiments (C and D) or are from one unbiased mass spectrometry screen (A and B). See also Figure S3.

### NSP14 SUMOylation inhibits SARS-CoV-2 ExoN activity

The cryo-EM structure of the SARS-CoV-2 NSP10-NSP14-RNA complex shows that K9 is within hydrogen bonding distance to the RNA backbone, while K200 is located at the interface that mediates binding of NSP14 to NSP10, which is crucial for efficient ExoN activity^5^ (**Figure 4E**).

We therefore posited that SUMOylation at these sites may inhibit ExoN enzymatic activity. To examine the effect of TRIM32-mediated SUMOylation on NSP14’s ExoN activity, we modified an established *in vitro* enzymatic activity assay^36^ by incubating NSP14, which was affinity-purified from mammalian cells co-expressing SUMO2, UBC9, and TRIM32 (WT, ΔRING, or mutSIM), with recombinant NSP10 and a coronaviral RNA substrate^36^ (**Figure 5A**). We found that NSP14 ExoN effectively degraded the RNA substrate in the absence of WT TRIM32; however, in the presence of WT TRIM32, ExoN activity was markedly suppressed. By contrast, NSP14 efficiently degraded the RNA in the presence of the TRIM32 ΔRING and mutSIM mutants (**Figure 5B**). Crucially, we confirmed in the same reaction samples that the ExoN activity inversely correlated with NSP14 SUMOylation by TRIM32 WT or mutants (**Figure S4A**). Notably, the ExoN-activity-dependent RNA degradation was confirmed in this assay by using a catalytically inactive NSP14 mutant (D90A/E92A)^7^ as control (**Figure S4B**).

**Figure 5.**
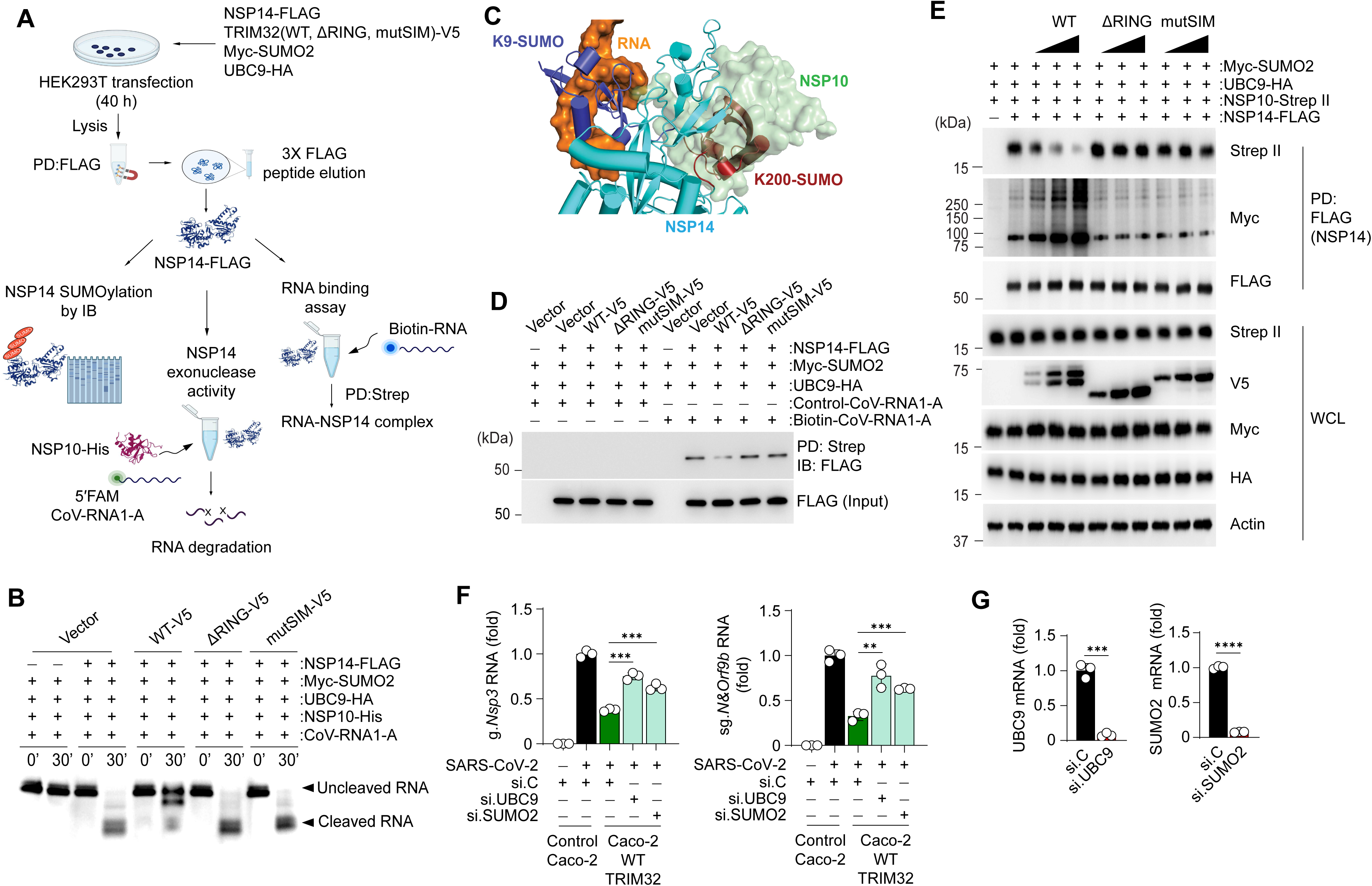
**ExoN SUMOylation by TRIM32 inhibits RNA and NSP10 binding, suppressing ExoN activity** (**A**) Schematic diagram of the *in vitro* exonuclease and RNA-binding assays to assess the effect of TRIM32-mediated SUMOylation on NSP14 ExoN activity and RNA-substrate binding. FLAG-tagged NSP14 or empty vector (control) was expressed in transiently transfected HEK293T cells together with either empty vector or TRIM32-V5 (WT, ΔRING or mutSIM) and Myc-SUMO2 and UBC9-HA, followed by affinity purification of NSP14-FLAG by PD: FLAG and subsequent elution using 3X-FLAG peptide. One portion of purified NSP14-FLAG was incubated with recombinant His-tagged NSP10 (rNSP10) and RNA substrate (CoV-RNA1-A) for the indicated times to analyze ExoN activity (lower center). Furthermore, eluted NSP14-FLAG was used for *in vitro* RNA binding assay with a biotinylated RNA substrate (CoV-RNA1-A) and subjected to PD: Strep and IB with anti-FLAG (lower right). The third portion of the eluted NSP14-FLAG fraction was analyzed by IB with anti-Myc to assess NSP14 SUMOylation (lower left). Parts of this figure were created using Biorender.com. (**B**) *In vitro* ExoN activity of purified SARS-CoV-2 NSP14 in the absence or presence of TRIM32 (WT or mutants) and the indicated SUMOylating machinery components, determined as described in (A). (**C**) Structural modeling of SARS-CoV-2 NSP14 with K9 and K200 SUMOylated. NSP10 (light green) and RNA (orange) from the NSP10-NSP14-RNA cryo-EM structure (PDB:7N0B) are depicted using surface representation showing steric clashes with SUMO molecules (K9-SUMO and K200-SUMO in NSP14 (cyan)). (**D**) RNA binding of SARS-CoV-2 NSP14-FLAG purified from mammalian cells co-expressing either empty vector or TRIM32 (WT or mutants) and the indicated SUMOylating machinery components and incubated with biotinylated CoV-RNA1-A (or control RNA), determined by PD: Strep and IB with anti-FLAG. (**E**) SARS-CoV-2 NSP14 binding to NSP10 in HEK293T cells that were co-transfected for 40 h with NSP14-FLAG, NSP10-Strep, UBC9-HA, Myc-SUMO2, and either empty vector or V5-tagged TRIM32 WT or mutants (ΔRING or mutSIM), determined by PD: FLAG and IB with anti-Strep. NSP14 SUMOylation was assessed in parallel by PD: FLAG and IB with anti-Myc. WCLs were probed by IB with anti-FLAG, anti-Strep, anti-V5, anti-Myc, anti-HA, and anti-Actin (loading control). (**F**) SARS-CoV-2 replication in stable Caco-2 cells that inducibly (by DOX treatment (2 µg/mL) for 24 h) expressed FLAG-tagged TRIM32 and were transfected for 40 h with si.C, si.UBC9, or si.SUMO2, followed by infection with SARS-CoV-2 (strain WA1; MOI 0.02). At 48 h post-infection, SARS-CoV-2 g.*Nsp3* and sg.*N&Orf9b* transcripts were determined by RT-qPCR. (**G**) Representative knockdown efficiency of endogenous UBC9 and SUMO2 for the experiment shown in (F), determined by RT-qPCR. Data (B, D, E, F, G) are representative of at least two independent experiments (mean ± s.d. of n = 3 biological replicates (F and G). **p < 0.01, ***p < 0.001, ****p < 0.0001 (Student’s *t* test). ns, not significant. See also Figure S4.

Since poly-SUMO2 chains are a high-molecular-weight ‘bulky’ moiety, we hypothesized that the SUMOylation of NSP14 may sterically impede its ability to bind NSP10 and/or RNA. In line with this concept, structural modeling of SUMOylated NSP14 suggested that the SUMO modifications at K9 and K200 sterically preclude RNA binding and NSP10 recruitment, respectively (**Figure 5C**). Indeed, functional assays showed that WT TRIM32 markedly diminished NSP14 binding to a SARS-CoV-2 RNA-substrate and to NSP10 (**Figures 5D and 5E**). By contrast, no such binding inhibition was observed in the presence of TRIM32 ΔRING or mutSIM (**Figures 5D and 5E**), strengthening the concept that SUMOylation of NSP14 by TRIM32 sterically inhibits these key interactions and thereby blocks ExoN activity.

To determine the physiological relevance of TRIM32-mediated SUMOylation for SARS-CoV-2 restriction, we determined the effect of enforced TRIM32 expression, with or without UBC9 or SUMO2 silencing, on SARS-CoV-2 replication (**Figures 5F and 5G**). As compared to cells expressing empty vector, inducible expression of WT TRIM32 potently restricted SARS-CoV-2 replication in cells that were co-transfected with non-targeting control siRNA (si.C). By contrast, we observed significantly diminished SARS-CoV-2 restriction by ectopic TRIM32 expression in cells depleted of UBC9 and SUMO2 (**Figures 5F and 5G**). These data reveal a novel SARS-CoV-2 restriction mechanism in which TRIM32 catalyzes SUMOylation of NSP14’s ExoN domain, which inhibits RNA-substrate binding and NSP10 recruitment, restraining ExoN activity.

### NSP14 SUMOylation by TRIM32 is broadly conserved in coronaviruses

Coronavirus NSP14 proteins, including their ExoN domains, are highly conserved among different coronaviruses (both alpha-and beta-coronaviruses) (**Figure S4C**). Therefore, we determined whether the NSP14 proteins of other coronaviruses, in particular SARS-CoV, MERS-CoV, HCoV-OC43, HCoV-229E, HCoV-NL63 and mouse hepatitis virus (MHV), also undergo SUMOylation by TRIM32. We found that these coronaviral NSP14 proteins were also robustly SUMOylated by TRIM32, where some NSP14s showed even higher SUMOylation levels than NSP14 from SARS-CoV-2 (**Figure 6A**). In contrast to WT TRIM32, the ΔRING mutant did not potentiate SUMOylation of the coronaviral NSP14 proteins, strengthening that SUMOylation is a direct effect of TRIM32’s RING E3 enzymatic activity (**Figure 6A**). In line with this, all the coronavirus NSP14 proteins tested efficiently interacted with V5-tagged TRIM32. By contrast, SARS-CoV-2 NSP10 (included as a negative control) did not bind TRIM32-V5 under the same conditions (**Figure 6B**). To determine whether TRIM32 inhibits other coronaviruses through its SUMOylation activity, we evaluated the effect of enforced TRIM32 expression on MHV replication in cells depleted of endogenous UBC9 or SUMO2. Cells stably expressing TRIM32 and transfected with si.C served as control (**Figures 6C and 6D**). This showed that TRIM32 markedly suppressed MHV replication in si.C-transfected cells; however, TRIM32-mediated MHV restriction was reversed by UBC9 or SUMO2 silencing. Collectively, these findings suggest that TRIM32-mediated SUMOylation of NSP14 is a conserved host restriction mechanism for a range of coronaviruses, both high-pathogenic and low-pathogenic ones.

**Figure 6.**
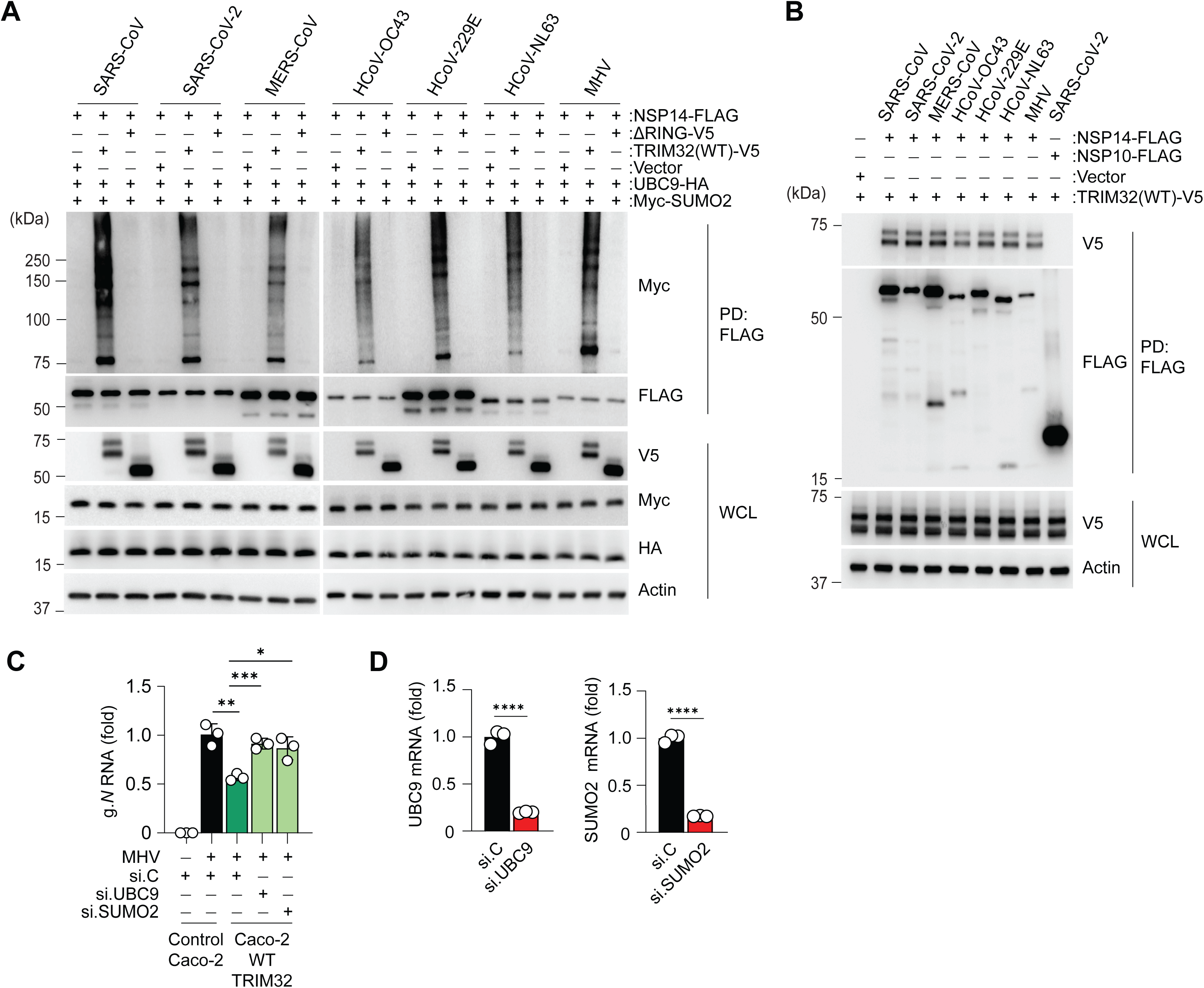
**NSP14 SUMOylation by TRIM32 is broadly conserved for coronaviruses** (**A**) SUMOylation of NSP14 from the indicated coronaviruses in HEK293T cells that were transiently transfected with plasmids expressing FLAG-tagged coronaviral NSP14s together with UBC9-HA and Myc-SUMO2 and either empty vector or V5-tagged TRIM32, assessed by PD: FLAG and IB with anti-Myc. WCLs were probed by IB with the indicated antibodies. (**B**) TRIM32 binding of the indicated coronaviral NSP14 proteins in HEK293T cells that were transiently transfected with plasmids expressing FLAG-tagged coronaviral NSP14s together with either empty vector or V5-tagged TRIM32, determined by PD: FLAG and IB with anti-V5. FLAG-tagged NSP10 (SARS-CoV-2) served as negative control. (**C**) MHV replication in stable Caco-2 cells that inducibly (by DOX treatment (2 µg/mL) for 24 h) expressed FLAG-tagged TRIM32 and were transfected for 40 h with si.C, si.UBC9 or si.SUMO2, followed by infection with MHV (strain A59; MOI 0.02) for 48 h, determined by RT-qPCR of *g.N* transcripts. (**D**) Representative knockdown efficiency of endogenous UBC9 and SUMO2 for the experiment in (C), determined by RT-qPCR. Data (A—D) are representative of at least two independent experiments (mean ± s.d. of n = 3 biological replicates (C and D). *p < 0.05, **p < 0.01, ***p < 0.001, ****p < 0.0001 (Student’s *t* test). See also Figure S4.

## DISCUSSION

The work from many groups has shown that TRIM proteins are important host-regulatory proteins negatively or positively influencing virus replication and disease outcome^10–13^. While the roles of many TRIM proteins in influenza, HIV, and herpesvirus infections have been extensively studied, little is still known about their relevance in modulating SARS-CoV-2 infection. The work presented herein comprehensively catalogued anti- and pro-viral TRIM proteins during SARS-CoV-2 infection by performing a human TRIM-wide RNAi screen. Our screen not only confirmed the important role of some TRIMs previously shown to modulate SARS-CoV-2 replication (*i.e.* TRIM7, TRIM21)^37,38^, but also revealed multiple TRIM proteins with previously unknown biological functions in SARS-CoV-2 infection (*i.e.* TRIM32, TRIM55, TRIM65, TRIM70). Furthermore, our data together with previous findings^39^ suggest that some TRIM proteins, such as TRIM6, can act anti-or pro-viral in a cell-type-specific manner. At the molecular level, while some of these TRIM proteins are known to modulate cytokine responses or autophagy (of note, most of them have not been studied in the context of SARS-CoV-2), future studies are needed to characterize the precise mechanisms by which these TRIM proteins regulate SARS-CoV-2 replication, including which substrate(s) are modified by their E3 ligase activities, or whether certain TRIM proteins perhaps influence SARS-CoV-2 replication in an E3 ligase-independent manner.

One of the top hits that emerged from our screen was TRIM32 that inhibited SARS-CoV-2 infection in a RING E3 ligase-dependent but IFN-independent manner. TRIM32 is known to restrict other RNA viruses, including IAV, VSV, and Newcastle Disease virus, where it either targets viral components for ubiquitination-dependent proteasomal degradation^40^ or potentiates antiviral type I IFN responses by catalyzing K63-linked polyubiquitin on innate signaling proteins^23^. Moreover, TRIM32 inhibits certain alphaviruses by blocking incoming virus after membrane fusion^41^, although the precise mechanism and substrate(s) in this context are elusive. The work presented herein discovered that the coronaviral NSP14 is a direct target of TRIM32, which blocks its ExoN enzymatic activity through direct SUMOylation, unveiling an unprecedented mechanism of TRIM-mediated virus restriction.

Our proteomics and biochemical characterization studies showed that TRIM32 catalyzes polymeric SUMO2 chains on NSP14 at two residues, K9 and K200, in the N-terminal ExoN domain. These residues are located at functional interfaces of NSP14. K9 locates to the ExoN substrate-binding pocket mediating viral RNA binding, while K200 lies at the interface responsible for NSP10 co-factor binding. Our functional analyses suggested that TRIM32-mediated SUMOylation sterically blocks NSP14 binding to both RNA and NSP10. As the NSP14 ExoN proofreading activity is known to critically contribute to SARS-CoV-2 resistance to nucleoside analogs inhibiting the viral RNA-dependent RNA polymerase^5,42,43^, it is tempting to speculate that agents to potentiate TRIM32’s ExoN-SUMOylation activity, or perhaps, more broadly, agents to boost the host’s SUMOylation machinery, may represent a new avenue for enhancing the efficacy of nucleoside analog antiviral drugs.

The NSP14 proteins of different coronaviruses share high sequence and structure similarity, and our data suggested that ExoN SUMOylation and virus restriction by TRIM32 are broadly conserved among coronaviruses. Determining whether K9 and K200, which are highly conserved (**Figure S4D**), are the major SUMOylation sites also in other coronaviral NSP14 proteins requires additional investigation. Moreover, future research is warranted to define the physiological relevance of TRIM32 in controlling coronavirus infection and pathogenesis in infected hosts. Another key question to be addressed is how SARS-CoV-2 (and other coronaviruses) modulate or antagonize TRIM32-mediated ExoN SUMOylation to maintain their genomic stability and to replicate efficiently.

Our study identified TRIM32, known to mediate ubiquitination events^23,40,44^, as a SUMO E3 ligase. Further investigation is needed to define the precise molecular architecture of the TRIM32-NSP14-UBC9 complex and to determine the mechanistic details of how SUMO2 is transferred by UBC9 onto NSP14’s ExoN in the presence of TRIM32. Moreover, the molecular details of how TRIM32 stabilizes growing poly-SUMO chains, and which structural elements in TRIM32 mediate this elongation, have yet to be determined. Similarly, the precise function of, and the specific reactions facilitated by, the proposed SIM motif in the TRIM32 NHL domain remain to be elucidated, and whether perhaps other motifs are required for TRIM32’s SUMOylation activity. Along these lines, it remains to be determined whether TRIM32 in complex with SUMO2, UBC9, and NSP14 also forms CCD-mediated antiparallel higher-order assemblies and whether these regulate TRIM32’s SUMOylation catalytic activity, alike previously shown for its ubiquitin ligase activity^45^. More broadly, our studies identifying TRIM32 as a SUMO E3 ligase provide a mandate for investigating other cellular substrates of its SUMOylation activity, both in other viral infections and in non-viral pathologies, such as hereditary muscular diseases (*i.e.* Limb-Girdle muscular dystrophy R8 and sarcotubular myopathy) which are associated with specific TRIM32 mutations^46,47^.

Taken together, this study unveils a host restriction mechanism of SARS-CoV-2 replication that targets the viral ExoN activity. Our work provides new insight into viral exonuclease regulation by the host SUMOylation machinery, which may spur new avenues for therapeutics design for coronavirus infection in humans.

## ACKNOWLEDGEMENTS

We greatly thank Bellinda Willard (Proteomics Core, Cleveland Clinic) for her support with the mass spectrometry analysis. We are also grateful to Gack lab members, in particular GuanQun Liu (now McGill University), for advice and technical support with the RNAi screen. This study was supported by grants from the U.S. National Institutes of Health (AI148534, AI087846 and AI169444 to M.U.G.).

## AUTHOR CONTRIBUTIONS

Study design and conceptualization, K.B., S.C. and M.U.G.; Methodology, K.B., S.C., C.C. and M.U.G.; Investigation, data analysis and visualization, K.B., S.C., C.C. and M.U.G.; Analysis of PDB structures: S.T, A.A.T. and S.K.O.; Writing – Original Draft, K.B., S.C., C.C. and M.U.G.; Funding Acquisition, S.K.O. and M.U.G.; Supervision, S.K.O. and M.U.G.

## DECLARATION OF INTEREST

The authors report no competing interests.

## SUPPLEMENTARY FIGURE LEGENDS

**Figure S1. Cell viability and knockdown efficiency analyses for the TRIM RNAi screen**

A. Cell cytotoxicity analysis for each sample of the TRIM RNAi screen shown in Figure 1B, assessed at 72 h post-infection by Hoechst 33342 staining and quantification of fluorescence intensity using a microplate reader. Values indicate percentage of live cells.
B. Representative knockdown efficiency of the indicated TRIM genes as well as control genes (*i.e.* LY6E and TMPRSS2) in Caco-2-hACE2 cells for the experiments shown in Figures 1E—H, determined by RT-qPCR. Values were normalized to cellular *HPRT1* and are presented compared to si.C, set to 1.
C. Representative knockdown efficiency of the indicated TRIM and control genes in Calu-3 cells for the experiments shown in Figures 1I—L, determined by RT-qPCR. Values were normalized and presented as in (B).

Data are representative of at least one screen (A) or two independent experiments (B and C) (mean ± s.d. of n = 3 technical (A) or biological replicates (B and C)). *p < 0.05, **p < 0.01, ***p < 0.001, ****p < 0.0001 (Student’s *t* test). ns, not significant.

See also Figure 1.

**Figure S2. SARS-CoV-2 replication in Caco-2 cells stably expressing TRIM32 WT or ΔRING**

A. Validation of stable Caco-2 cells inducibly expressing TRIM32 WT or ΔRING. Expression of FLAG-tagged TRIM32 WT or ΔRING mutant in Caco-2 cells engineered to inducibly express those proteins, determined in the WCLs by IB with anti-FLAG after incubation of cells with 1 µg/mL DOX for 24 h. Caco-2 cells stably expressing empty vector served as control.
B. SARS-CoV-2 replication kinetics over a 72-h period in stable Caco-2 cells inducibly (via 1 µg/mL DOX treatment for 24 h) expressing either TRIM32 WT or ΔRING, or in control (vector-expressing) cells, that were either mock-treated or infected with EGFP-rSARS-CoV-2 (strain K49; MOI 0.05), determined by measuring RFU for EGFP. Values were auto-scaled to the average values of mock-infected wells at 0 h, set to 200.

(**C** and **D**) SARS-CoV-2 g.*Nsp3* (C) and sg.*N&Orf9b* (D) transcripts in stable Caco-2 cells inducibly (via 1 µg/mL DOX treatment for 24 h) expressing either TRIM32 WT or ΔRING, or in control (vector-expressing) cells, that were either mock-treated (−) or infected with SARS-CoV-2 (strain WA1; MOI 0.05) for 48 h, determined by RT-qPCR.

A. Protein abundance of SARS-CoV-2 spike (S) and nucleocapsid (N) in stable Caco-2 cells expressing TRIM32 WT or ΔRING, or control cells, that were treated and infected as in (C), determined in the WCLs by IB with anti-S and anti-N.

(**F** and **G**) *IFIT2* (F) and *ISG15* (G) transcripts in control (vector-expressing) or stable Caco-2 cells inducibly expressing TRIM32 WT or ΔRING that were treated with DOX as in (C) and then treated as indicated with anti-IFNAR2 antibody (2 µg/mL) for 2 h and subsequently either mock-infected (−) or infected with SARS-CoV-2 (strain WA1; MOI 0.05) for 48 h, determined by RT-qPCR.

(**H**) Virus replication in stable Caco-2 cells expressing TRIM32 WT or ΔRING, or in control cells, treated with DOX and anti-IFNAR2 as in (F) and then mock-infected or infected with EGFP-rSARS-CoV-2 (strain K49; MOI 0.05) for 48 h, determined by assessing EGFP using fluorescence microscopy analysis.

Data (A—H) are representative of at least two independent experiments (mean ± s.d. of n = 3 biological replicates (B—D, F—G)). **p < 0.01, ***p < 0.001 (Student’s *t* test).

See also Figure 2.

**Figure S3. TRIM32 colocalization to viral RTCs, its effect on NSP14 ubiquitination, and its UBC9-binding ability**

(**A** and **B**) Co-localization of endogenous TRIM32 (green) with SARS-CoV-2 dsRNA (red) in A549-hACE2 (A) and Caco-2-hACE2 (B) cells that were either mock-treated or infected with SARS-CoV-2 (strain WA1; MOI 1) for 24 h, assessed by immunostaining with anti-TRIM32 and anti-dsRNA and confocal laser scanning microscopy. Nuclei, DAPI (blue). Scale bar, 10 μm.

Binding of V5-tagged TRIM32 to the indicated FLAG-tagged SARS-CoV-2 proteins in HEK293T cells that were transiently co-transfected for 40 h to express those proteins, or transfected with empty vector, determined by pull-down with anti-FLAG (PD: FLAG) and IB with anti-V5. WCLs were probed with anti-V5 and anti-Actin (loading control).
AlphaFold model of full-length TRIM32 in cartoon representation, with RING (yellow), BB (cyan), CC (pink), and NHL (green) domains shown. Zinc ions are depicted as grey spheres.
Ubiquitination of FLAG-tagged NSP14 (SARS-CoV-2) in HEK293T cells that were co-transfected for 40 h with V5-tagged TRIM32 WT or mutants, determined by PD: FLAG and IB with anti-pan-ubiquitin (Ub). WCLs were probed by IB with anti-V5, anti-pan-Ub, and anti-Actin (loading control).
Binding of HA-tagged UBC9 to V5-tagged TRIM32 WT or mutants in transiently transfected HEK293T cells, determined by PD: V5 and IB with anti-HA. WCLs were probed with anti-HA and anti-Actin (loading control).

Data (A—C, E and F) are representative of at least two independent experiments. See also Figure 4.

**Figure S4. ExoN activity of SARS-CoV-2 NSP14 WT and mutant, and alignment of the coronaviral NSP14 regions that contain K9 and K200**

A. SUMOylation of NSP14-FLAG by V5-tagged TRIM32 WT or mutants in the ExoN reaction samples for the experiment shown in Figure 5B, determined by PD: FLAG and IB with anti-Myc.
B. *In vitro* ExoN activity of FLAG-tagged NSP14 (SARS-CoV-2) WT or D90A/E92A mutant that were affinity-purified from transiently transfected HEK293T cells and then incubated with recombinant NSP10 (His tagged) and CoV-RNA1-A for the indicated times.
C. Heatmap representation of the protein sequence identity (in %) of coronaviral NSP14 ExoN domains determined by ClustalW.
D. Alignments of the NSP14 protein regions containing K9 and K200 from the indicated human coronaviruses, determined by ClustalW. Asterisks “*” indicate positions which have a single, fully conserved residue; colons “:” illustrate conservation between groups of strongly similar properties; periods “.” indicate conservation between groups of weakly similar properties.

Data (A and B) are representative of at least two independent experiments. See also Figures 5 and 6.

## STAR METHODS

### RESOURCE AVAILABILITY

#### Lead contact

Further information and requests for resources and reagents should be directed to and will be fulfilled by the lead contact, Michaela U. Gack.

#### Materials availability

All unique/stable reagents and materials generated in this study are available from the lead contact (M.U.G.) with a completed materials transfer agreement.

#### Data and code availability

- The datasets generated and/or analyzed during this study are either included in the paper and/or are available from the lead contact upon request.
- This paper does not report original code.
- Any additional information required to reanalyze the data reported in this work paper is available from the lead contact upon request.

#### EXPERIMENTAL MODEL AND SUBJECT DETAILS

##### Cells

HEK293T (human embryonic kidney cells) were purchased from ATCC and maintained in Dulbecco’s modified Eagle’s medium (DMEM) supplemented with 10% (v/v) fetal bovine serum (FBS), 1 mM sodium pyruvate, and 1% (v/v) of penicillin–streptomycin. Vero E6-TMPRSS2 cells (African green monkey kidney epithelial cells) were cultured in Dulbecco’s Modified Eagle’s Medium (DMEM) supplemented with 10% (v/v) FBS and 1 mM sodium pyruvate, 1% (v/v) of penicillin–streptomycin, and 30 µg/mL Blasticidin. Calu-3 (human lung adenocarcinoma cells) were cultured in Eagle’s Minimum Essential Medium (EMEM) supplemented with 10% (v/v) FBS and 1% (v/v) penicillin-streptomycin. Caco-2 (colorectal carcinoma cells), Caco-2-hACE2, and Caco-2 *TRIM32*-KO and Caco-2 cells inducibly expressing TRIM32 (WT or mutants) were cultured in Essential Medium (EMEM) supplemented with 20% (v/v) FBS and 1% (v/v) penicillin-streptomycin. All cell cultures were maintained at 37 °C in a humidified 5% CO_2_ incubator.

Purchased cell lines were authenticated by the respective vendors and were not validated further in our laboratory. KO cell lines and cells stably expressing TRIM32 were validated by confirming knockout or expression of the target protein, respectively. All cell lines used in our study were regularly checked for potential mycoplasma contamination by PCR.

##### Viruses

Origins and strains of the viruses used in the current study are described in the Key Resources Table. SARS-CoV-2 (strain 2019-nCoV/USA_WA1 2020) was propagated in Vero E6-hACE2 cells using published protocols^48^. EGFP-rSARS-CoV-2 was propagated in Vero E6-TMPRSS2 cells and has been described previously^49^. The methods used to assess viral titers are described in detail below. All work relating to live SARS-CoV-2, and RNA derived from it, was conducted in the BSL-3 facility of the Cleveland Clinic Florida Research and Innovation Center (CC-FRIC). All work conducted in this study was approved by the Institutional Biosafety Committee of CC-FRIC and in accordance with U.S. National Institutes of Health guidelines.

##### Plasmids

Plasmids expressing human TRIM32 WT and its mutants ΔRING, RING-BB, BB-CCD, and NHL were cloned into the pcDNA3.1–V5 vector using the restriction sites NheI and KpnI. TRIM32 C39S and mutSIM were generated by site-directed mutagenesis using TRIM32 WT as template. FLAG-tagged SARS-CoV-2 NSP2, NSP10, NSP14, and NSP16 were kindly provided by J. U. Jung (Cleveland Clinic, Ohio). Single (K9R, K200R) and double (K9R/K200R, D90A/E92A) mutants of SARS-CoV-2 NSP14 were generated by site-directed mutagenesis using FLAG-NSP14 as template. NSP10-Strep-Tag II was described previously^50^. The plasmids Myc-SUMO1, Myc-SUMO2, Myc-SUMO3, and UBC-9-HA were kindly provided by F. Full (University of Freiburg). All constructs were validated by Sanger sequencing (Azenta Life Sciences) or Nanopore sequencing (Plasmidsaurus). Transfections were performed using linear polyethylenimine (1 mg/mL solution in 20 mM Tris pH 6.8) or Lipofectamine 2000 following the manufacturer’s instructions.

##### Other reagents

Protease inhibitor cocktail (Sigma-Aldrich #P2714) was added at a concentration of 1:50 to cell lysates for pull down and immunoprecipitation assays. Monoclonal IFNAR2-neutralizing antibody (1:250, clone MMHAR-2) was obtained from PBL Assay Science. FLAG M2 and Protein G Dynabeads were obtained from Millipore Sigma and Invitrogen, respectively.

#### METHOD DETAILS

##### Cell lysis, immunoblot analysis and antibodies

Indicated cell lines or primary cells were lysed in Nonidet P-40 (NP-40) buffer (50 mM HEPES [pH 7.4], 150 mM NaCl, 1% (v/v) NP-40, 1 mM EDTA, 1:50 protease inhibitor cocktail) or RIPA buffer (Thermo Scientific, Cat #89901), and lysates were centrifuged at 21,000 x g for 20 min at 4°C. For pull-down and immunoprecipitation assays, cell lysates were incubated with anti-FLAG or anti-V5 conjugated beads for 4-6 h or anti-TRIM32 or anti-SUMO2/3 for 4 h, then washed thoroughly with NP-40 or RIPA buffer. SUMOylation immunoblot samples were subjected to a modified protocol. Briefly, cells transfected with FLAG-NSP14 (with or without HA-UBC9 and Myc-SUMO1/2/3) were lysed using a modified RIPA buffer (50 mM Tris-HCl [pH 7.5], 150 mM NaCl, 1% (v/v) NP-40, 2% SDS, 0.25% sodium deoxycholate, 1 mM EDTA, 10 mM NEM) followed by boiling at 95 °C and sonication. The lysates were then diluted 10-fold using the modified RIPA buffer to achieve a final SDS concentration at 0.2%, followed by centrifugation at 20,000 x g for 20 minutes at 4 °C to clear the samples of debris. WCLs were mixed with Laemmli SDS sample buffer and heated at 95 °C for 5 min.

SDS-PAGE and IB analyses were carried out using previously published protocols^48,51,52^. Briefly, membranes were blocked in 5% (w/v) non-fat dry milk in PBS-T (0.05% (v/v) Tween-20 in PBS) for 1 h at room temperature, followed by primary antibody incubation overnight at 4°C. Following primary antibody incubation, blots were probed with HRP-conjugated secondary antibody for 1 h at room temperature. Primary antibodies were used for IB at a dilution of 1:1000 (unless otherwise indicated) and include: anti-TRIM32, anti-SARS-CoV-2 spike (S), anti-SARS-CoV-2 nucleocapsid (N), anti-SARS-CoV-2 NSP14, anti-FLAG, anti-V5, anti-HA, anti-Myc, anti-Strep-Tag II, anti-pan-Ub, anti-SUMO1, anti-SUMO2/3 and anti-β-actin (1:5000).

##### Virus infection and titer analysis

Virus infection experiments and titer analysis were performed as described previously^48^. Briefly, cells were incubated with SARS-CoV-2-containing inoculum diluted in DMEM or EMEM (2% FBS) for 1-2 h. Inoculum was removed and replaced with the respective complete growth medium, and cells were incubated for the indicated times. Viral titers were determined in Vero E6-TMPRSS2 cells by plaque assay following protocols described previously^49^. Plaques were counted and reported as plaque forming unit/mL (PFU/mL) or focus forming unit/mL (FFU/mL), calculated as number of (plaques or foci / well) x (dilution factor) / (infection volume).

##### RNA purification and qRT-PCR

Total cellular RNA was purified using the E.Z.N.A. HP Total RNA extraction kit, per the manufacturer’s instructions. Equal amounts of RNA (25–100 ng) were used in a one-step qRT-PCR reaction using the SuperScript III Platinum One-Step qRT-PCR kit with ROX and commercially-available predesigned FAM reporter dye primers for the analyzed transcripts using a QuantStudio 6 Pro Real-Time PCR Machine (Applied Biosystems). Relative mRNA expression was normalized to that of *HPRT1*. The comparative CT method (ΔΔCT) was used to measure the transcript levels of each target gene.

##### TRIM-specific siRNA screen and other siRNA knockdown experiments

A custom TRIM siRNA library was purchased from Horizon Discovery (Cherry-Pick Custom Library Tool) for the TRIM RNAi screen. The TRIM-specific siRNAs as well as the other siRNAs used in our study are described in the Key Resources Table. Transient gene silencing was performed using 80 nM of gene-specific siGENOME SMARTpool siRNAs using Lipofectamine RNAiMax transfection reagent, according to the manufacturer’s instructions. Non-targeting siRNAs (siGENOME non-targeting siRNA Pool #2) were used as a control. Knockdown efficiency of specific genes was determined by RT-qPCR at the indicated times using pre-designed primers (IDT) indicated in previous sections.

For the TRIM RNAi screen, Caco-2-hACE2 cells were seeded into 96-well white-clear bottom plates (Nunc) (approximately 2.5 x 10^4^ cells per well) and reverse-transfected with 80 nM of TRIM gene-specific siGENOME SMARTpool siRNAs using RNAiMax transfection reagent according to the manufacturer’s instructions. 40 h later, cells were infected with EGFP-rSARS-CoV-2 in FluoroBrite DMEM (Gibco) containing 2% (v/v) FBS, 1 mM sodium pyruvate, 1× NEAA, and 10 mM HEPES. Real-time EGFP fluorescence measurement was performed at 1-h intervals over a 72-h period in a temperature-and CO_2_-controlled Synergy Neo2 Hybrid Multi-Mode Reader (BioTek) using predefined monochromator/bandwidth settings for EGFP (excitation: 479/20; emission: 520/20). Gain values were automatically scaled to the average of mock-infected wells at 0 h, set to 200.

##### Generation of CRISPR KO and stable cells

Caco-2 *TRIM32*-KO cells were generated by CRISPR-Cas9-mediated genome editing using a TRIM32 CRISPR/Cas9 KO plasmid (sc-402731; Santa Cruz). Briefly, Caco-2 cells were transiently transfected with the TRIM32 CRISPR-Cas9 KO plasmid using Lipofectamine 3000. 40 h later, transfected cells that express GFP were subjected to fluorescence-activated cell sorting (FACS) analysis to sort single-cell clones. Knockout of TRIM32 expression was confirmed by RT-qPCR and IB to confirm the absence of TRIM32 protein expression.

Stable Caco-2 cells inducibly expressing TRIM32 WT or ΔRING (cloned into pGLVX-TetOne-vector with a C-terminal FLAG tag) were generated by lentiviral transduction, followed by selection with 5 µg/mL puromycin. Expression of TRIM32 WT and ΔRING mutant in these cells was confirmed after treatment with DOX (for the indicated times) by IB analysis of WCLs using an anti-FLAG antibody.

##### Identification of NSP14 SUMOylation sites by mass spectrometry

To identify the SUMOylation sites in SARS-CoV-2 NSP14, twenty-five 10 cm-dishes of HEK293T cells (∼1 x 10^8^ cells per dish) were transfected with FLAG-tagged NSP14, TRIM32 (WT)-V5, Myc-SUMO2 T91R, and UBC9-HA. At 40 h post-transfection, cells were harvested, washed with PBS, and lysed in 1% (v/v) NP-40 lysis buffer containing 10 mM NEM and 1 × protease inhibitor cocktail. Cell lysates were clarified by centrifugation at 21,000 x g for 20 min at 4 °C, and supernatant was incubated with anti-FLAG M2 magnetic beads at 4°C for 4 h. Pulldown samples were washed harshly with NP-40 lysis buffer. Samples were run on a 4-12% Bis-Tris commercial gel, stained with Coomassie blue, and bands corresponding to NSP14 were excised and subjected to trypsin digestion followed by LC-MS/MS at the Cleveland Clinic Lerner Research Institute Proteomics & Metabolomics Core to identify sites of post-translational modifications of NSP14.

##### Confocal microscopy

Cells were grown in 8-chamber slides (∼30,000 cells per well) and infected with virus as indicated. The cells were fixed with 4% (w/v) paraformaldehyde and permeabilized with 0.2% (v/v) Triton X-100 (in PBS). Next, cells were blocked in a PBS solution containing 2% (w/v) BSA and 0.05% (v/v) Triton-X-100 for 1 h at room temperature. After washing with PBS, cells were incubated on a rocking platform with the primary antibodies, specifically anti-TRIM32 (1:200) and dsRNA (J2) (1:500), diluted in PBS containing 1% (w/v) BSA) at 4 °C overnight. The next day, cells were washed three times with PBS containing 0.25% (v/v) Tween-20 and then incubated with secondary antibodies conjugated to Alexa Fluor 488 (1:500) and Alexa Fluor 647 (1:500). Cells were washed three times with PBS containing 0.25% (v/v) Tween-20, mounted in Prolong Gold Antifade mountant after DAPI staining, and imaged using a Leica Stellaris 8 confocal microscope.

##### ExoN activity assay

FLAG-tagged NSP14 protein was affinity-purified from HEK293T cells following a published FLAG-protein purification protocol^52^ with slight modifications. In brief, twenty 10 cm-dishes of HEK293T cells (∼1 x 10^8^ cells per dish) were transfected with FLAG-tagged NSP14 (or 3 X FLAG empty vector as a control) together with either empty vector or V5-tagged TRIM32 (WT, ΔRING, or mutSIM) along with Myc-SUMO2 and UBC9-HA. Forty hours later, the cells were washed with PBS and lysed in NP-40 buffer (50 mM HEPES [pH 7.4], 150 mM NaCl, 1% (v/v) NP-40, 1 mM EDTA, 10mM NEM, 1:50 protease inhibitor cocktail) on ice for 20 min. The samples were centrifuged at 15,000 rpm for 25 min at 4 °C. Cleared lysates were incubated with FLAG M2 agarose beads for 4 h at 4 °C. The samples were washed vigorously 7-10 times by vortexing at high speed. NSP14-FLAG protein was eluted from beads by incubation with 100 μg/mL 3X FLAG peptide in TBS (10 mM Tris-HCl, 150 mM NaCl [pH 7.4]) for 1 h at 4 °C at 650 rpm. The eluted protein concentration was measured by BCA assay. One portion of the eluted fraction was used to determine SUMOylation of NSP14 by IB analysis; the other portion of eluted NSP14-FLAG was used for *in vitro* exonuclease assay by incubating it with recombinant NSP10-His protein and RNA substrate (*i.e.* CoV-RNA1-A) using a published protocol^36^. Briefly, FAM-labelled RNA substrate (CoV-RNA1-A 5′ FAM-AAAAAAAAAAACGCGUAGUUUUCUACGCG 3′) was custom-synthesized (IDT). Equal amounts (1 μM) of FLAG-eluted NSP14 protein were mixed with 2 μM of recombinant NSP10 (ratio 1:2) in reaction buffer [20 mM HEPES (pH 7.5),100 mM NaCl, 5% glycerol, 10 mM MgCl2, 5 mM β-ME] on ice for 5 min. The protein complex was then incubated with 2 μM FAM-labeled RNA substrate and incubated at 37 °C. The reaction was stopped at the indicated times by adding 2x RNA loading dye, which contains formamide. The samples were then boiled for 5 min at 95 °C and resolved on a 15% TBE-Urea polyacrylamide gel at a constant power of 100 V for 60 min. RNA degradation was visualized with the Amersham Image Quant 800.

##### In vitro RNA binding assay

Biotin-RNA pulldown assay was performed using a published protocol with modifications^53^. To analyze RNA binding of SARS-CoV-2 NSP14, MgCl_2_ was substituted with CaCl_2_ in the RNA-protein binding buffer to prevent RNA cleavage by NSP14^5^. Briefly, biotinylated and control RNA oligonucleotides were custom-synthesized (IDT). 0.1 μM of biotinylated and control CoV-RNA1-A were immobilized onto 100 μL of streptavidin magnetic beads (Dynabeads™ M-280 Streptavidin Invitrogen™11206D) at 4 °C for 4 h. RNA-conjugated streptavidin beads were then incubated with equal amounts of eluted NSP14-FLAG protein in RNA-protein binding buffer (20 mM HEPES [pH 7.5], 100 mM NaCl, 4 mM CaCl_2_, 5 mM β-ME, 5% glycerol) at 4 °C for 8 h. Beads containing RNA-NSP14 complexes were washed three times with washing buffer (20 mM HEPES [pH 7.5], 200 mM NaCl, 0.05% NP-40, 1 mM EDTA). RNA-NSP14 complexes were eluted in 2X Laemmli sample buffer. The samples were then boiled for 5 min at 95 °C, resolved on a 10% Bis-Tris gel, and subjected to immunoblotting with anti-FLAG.

##### Sequence alignments and structural modeling

Primary sequence alignments of coronaviral NSP14 proteins were performed using Clustal Omega. Illustration of the published SARS-CoV-2 NSP10-NSP14-RNA structure (PDB: 7N0B) was performed in PyMOL. The modelling of the full-length TRIM32 structure was performed using AlphaFold.

##### QUANTIFICATION AND STATISTICAL ANALYSIS

Data are presented as mean ± s.d. and analyzed using GraphPad Prism software (version 10). An unpaired two-tailed Student’s *t*-test was used. Pre-specified effect sizes were not assumed and generally three biological replicates (n) for each condition were used, unless indicated otherwise. Data were reproduced in independent experiments as indicated in the legend for each figure.

## KEY RESOURCES TABLE

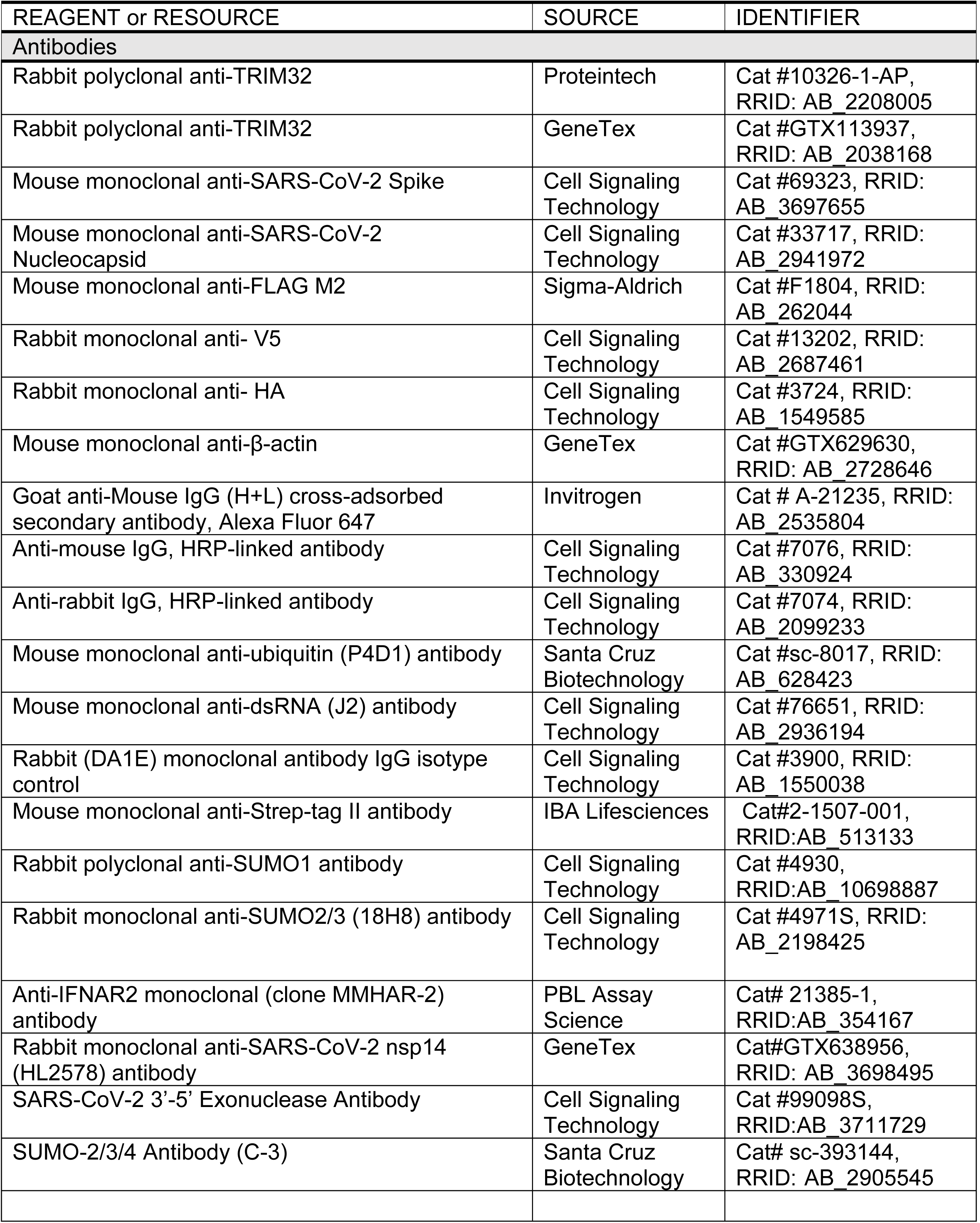
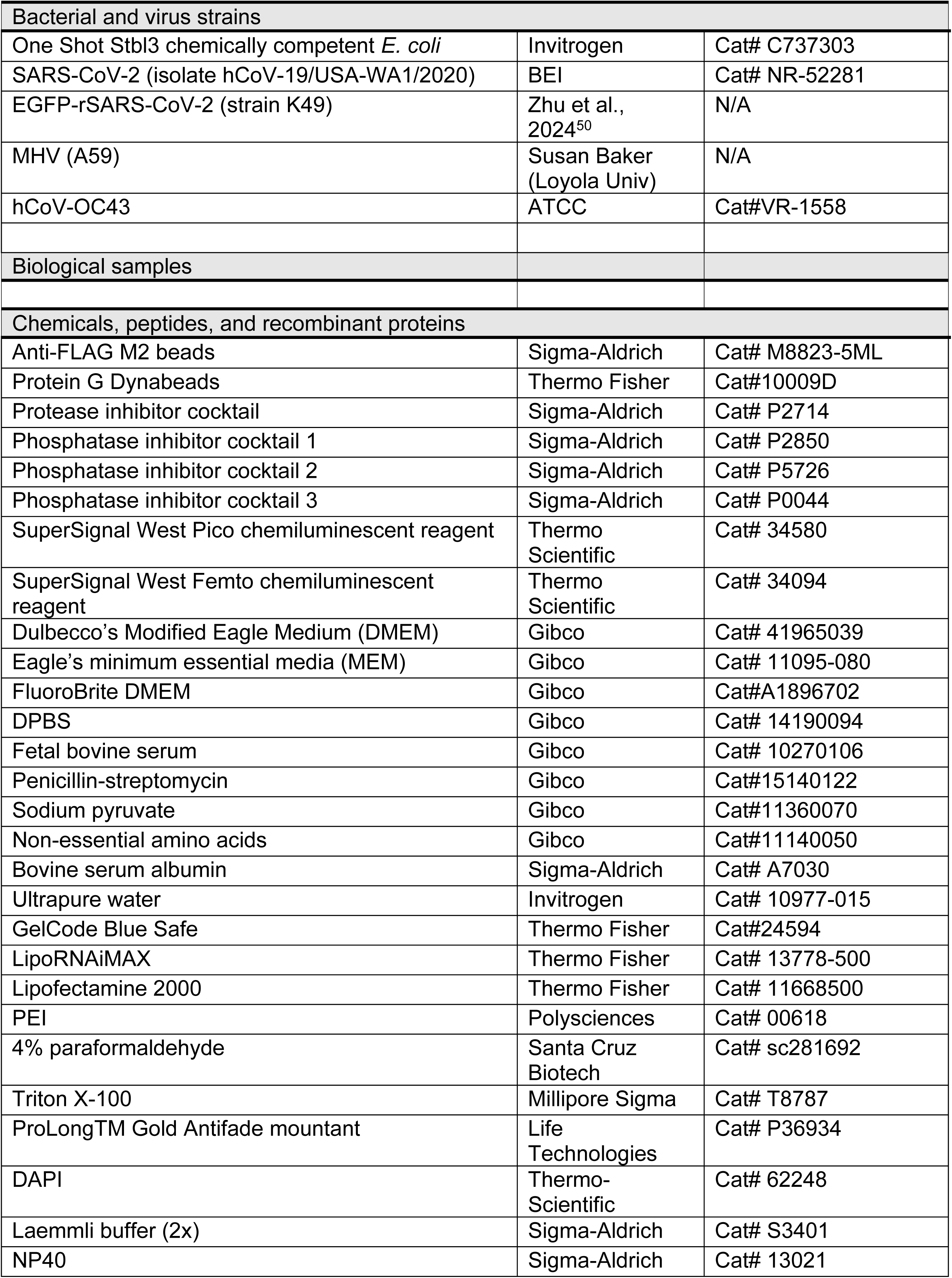
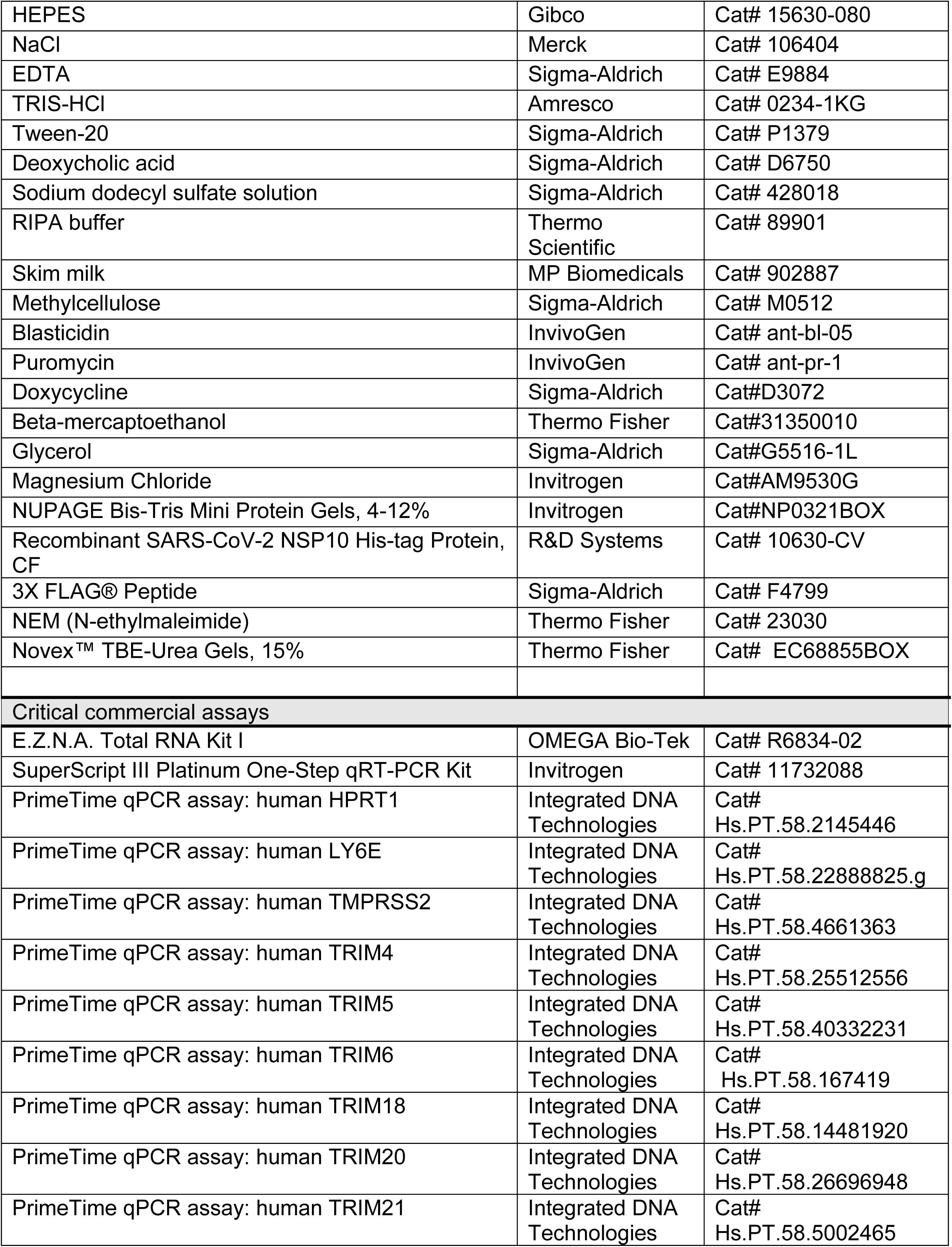
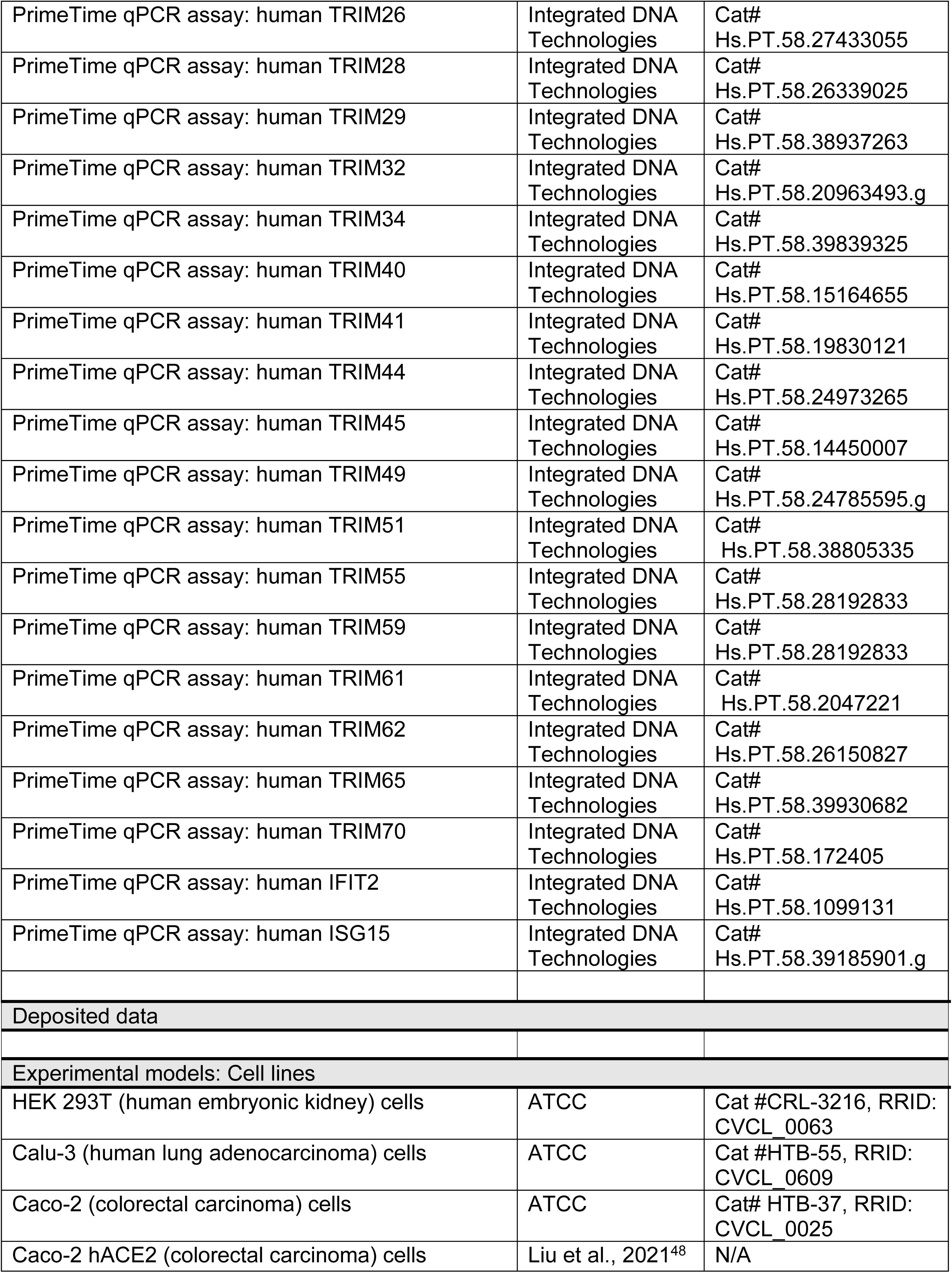
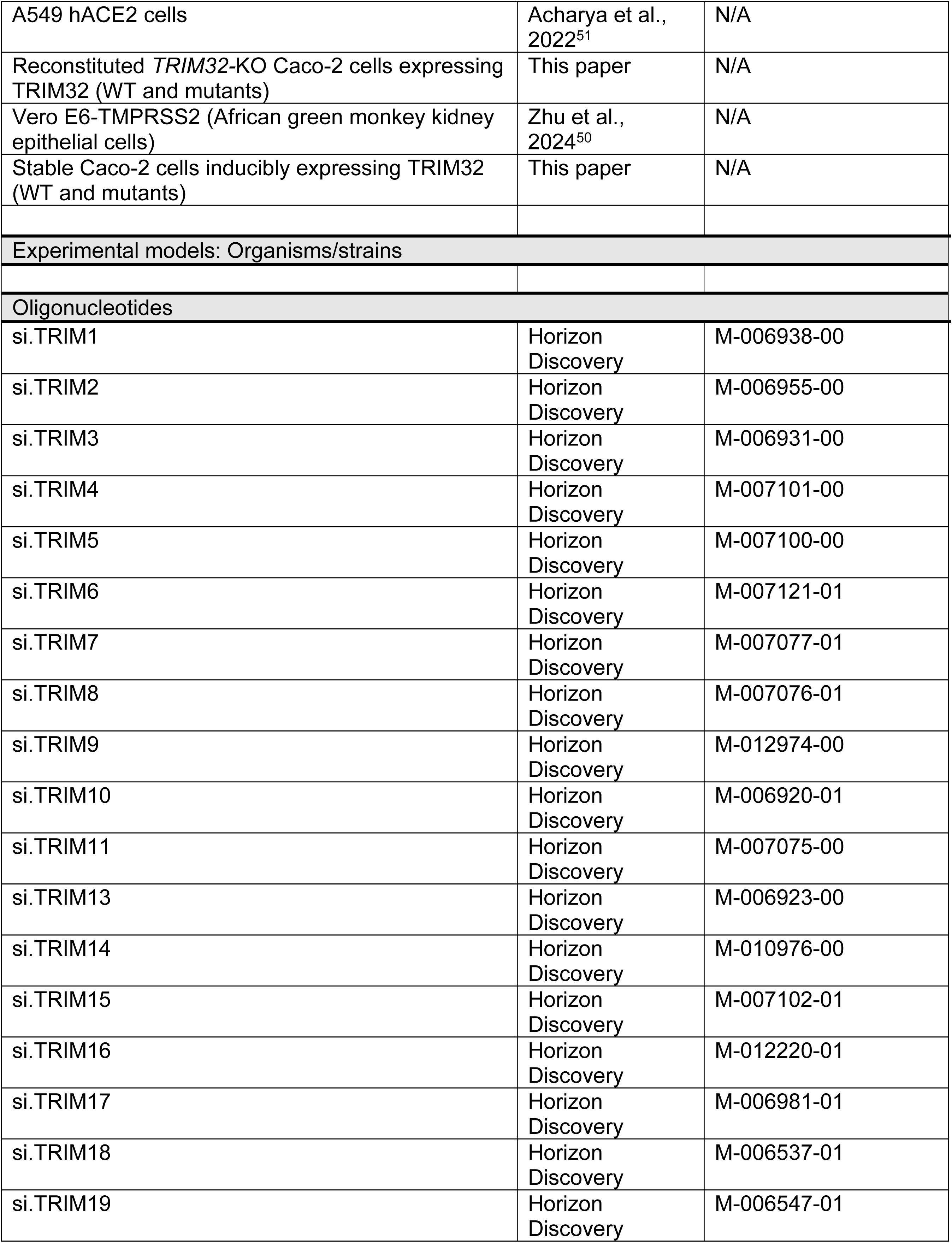
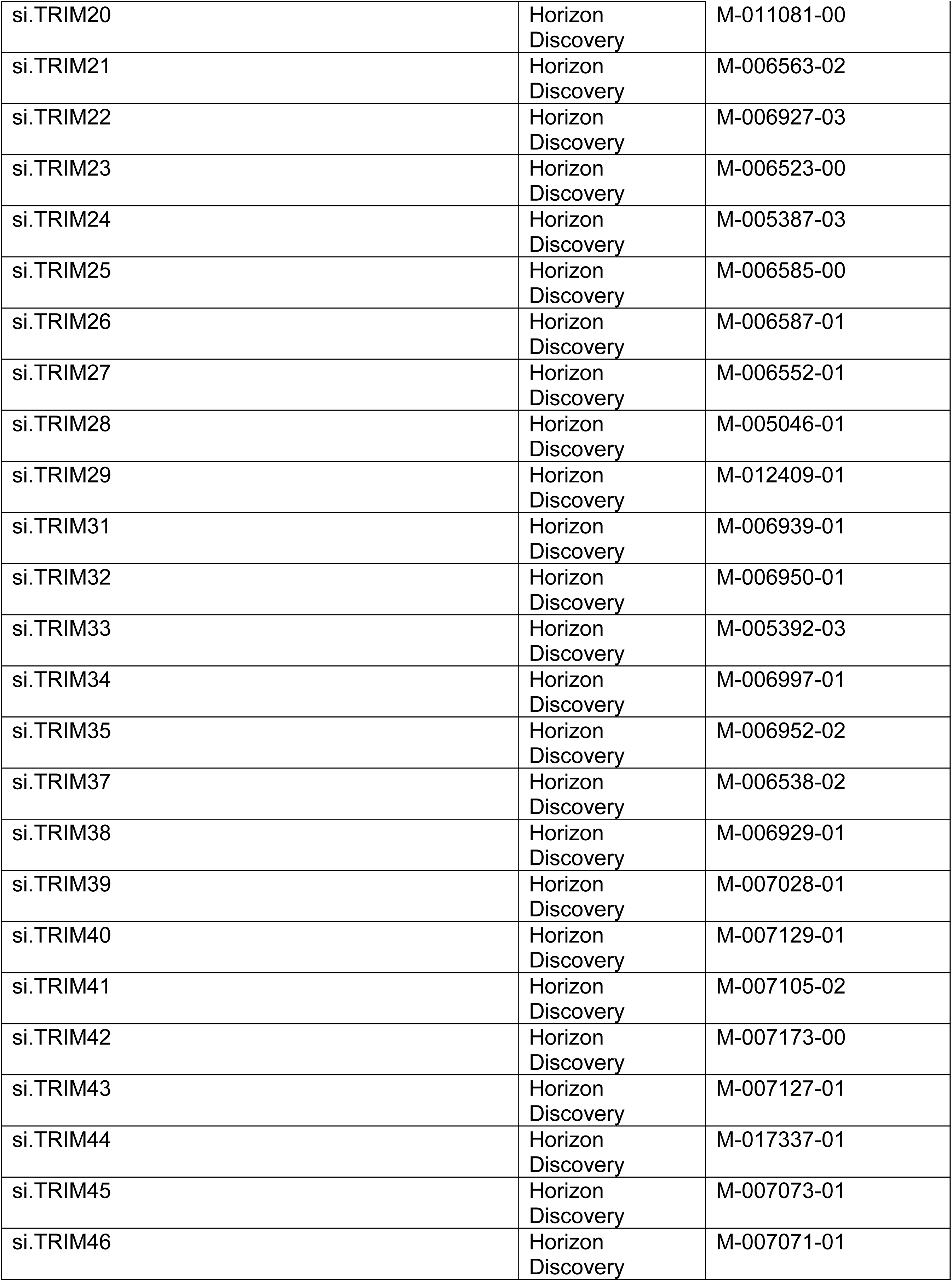
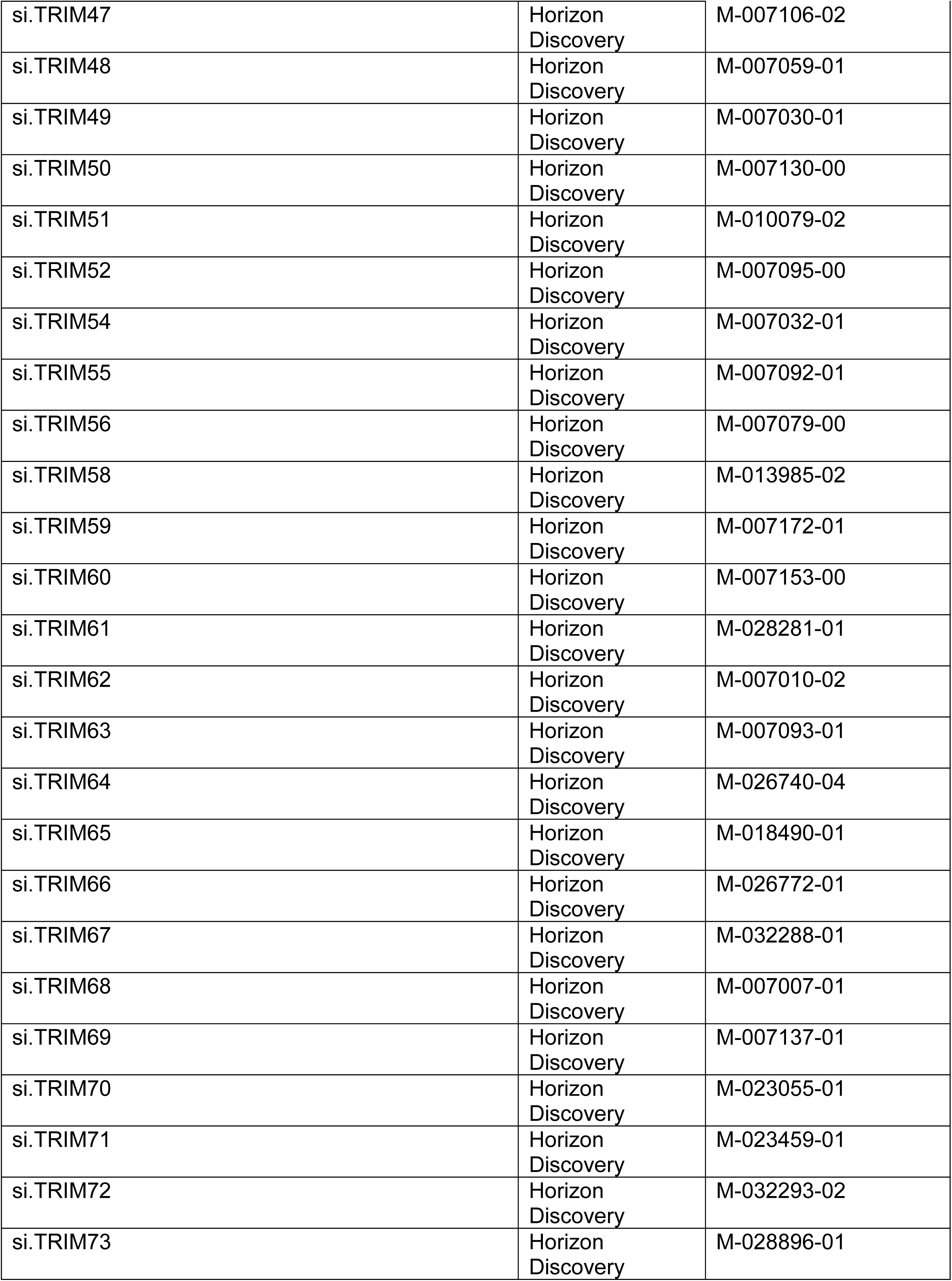
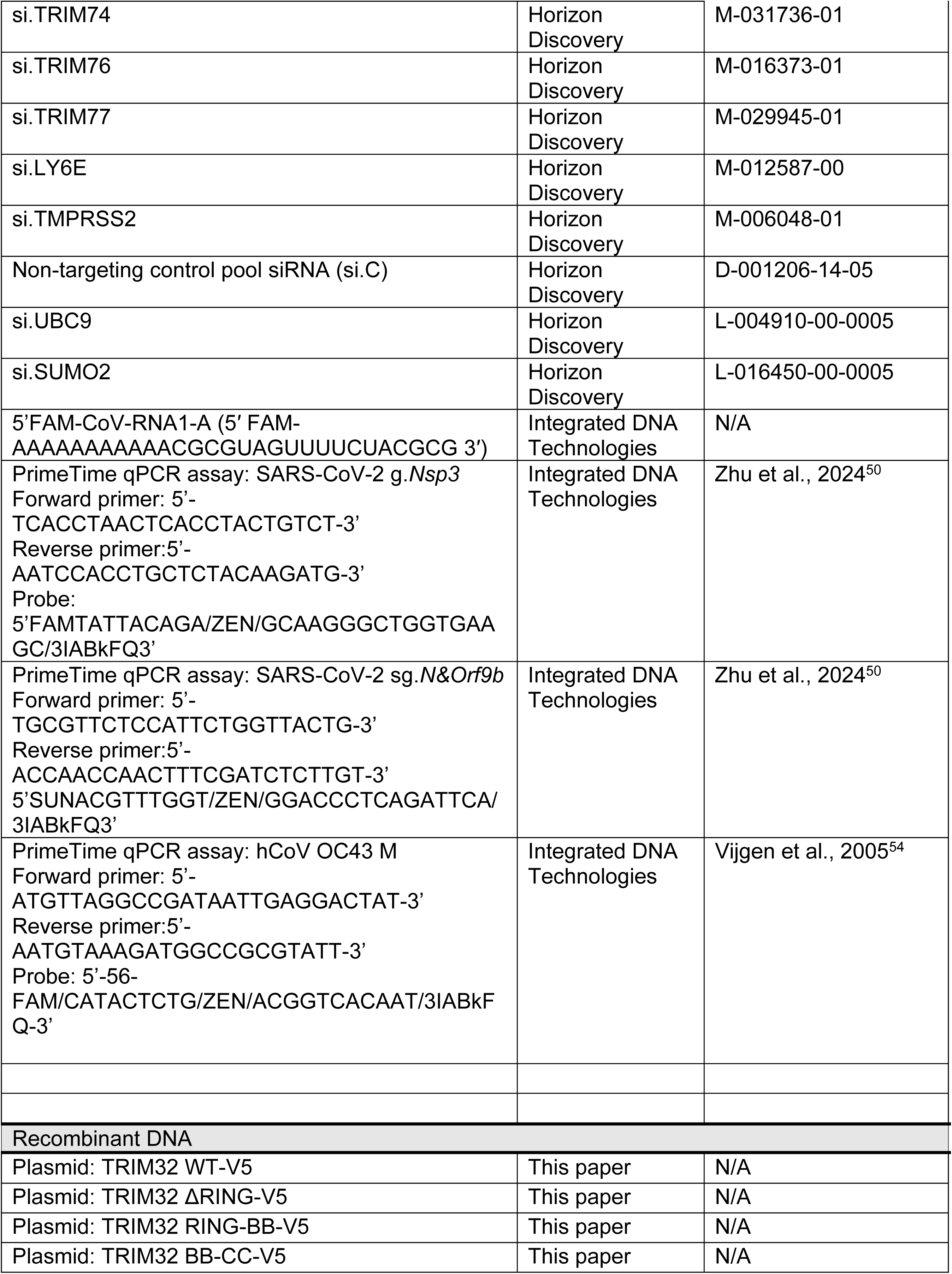
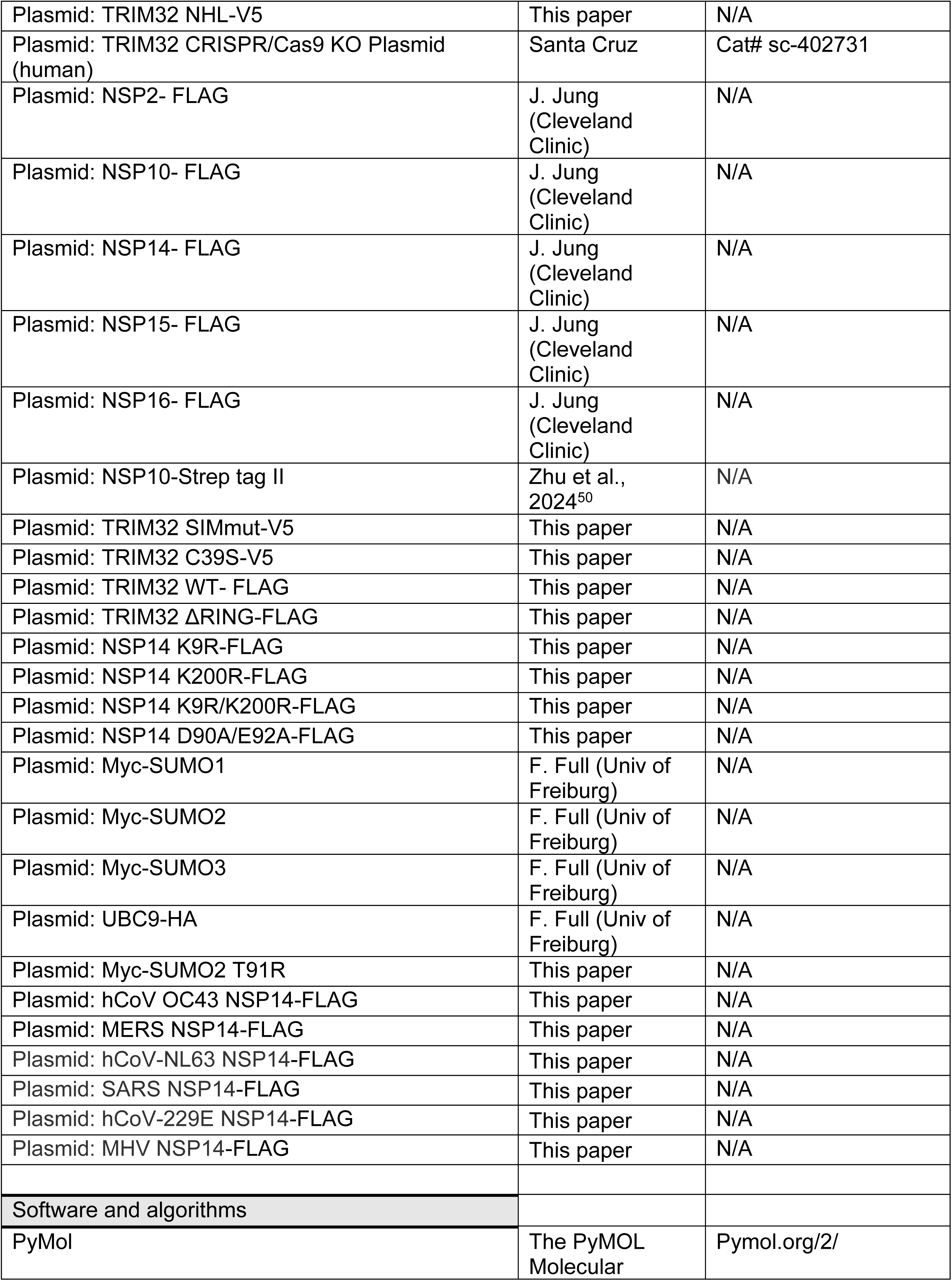
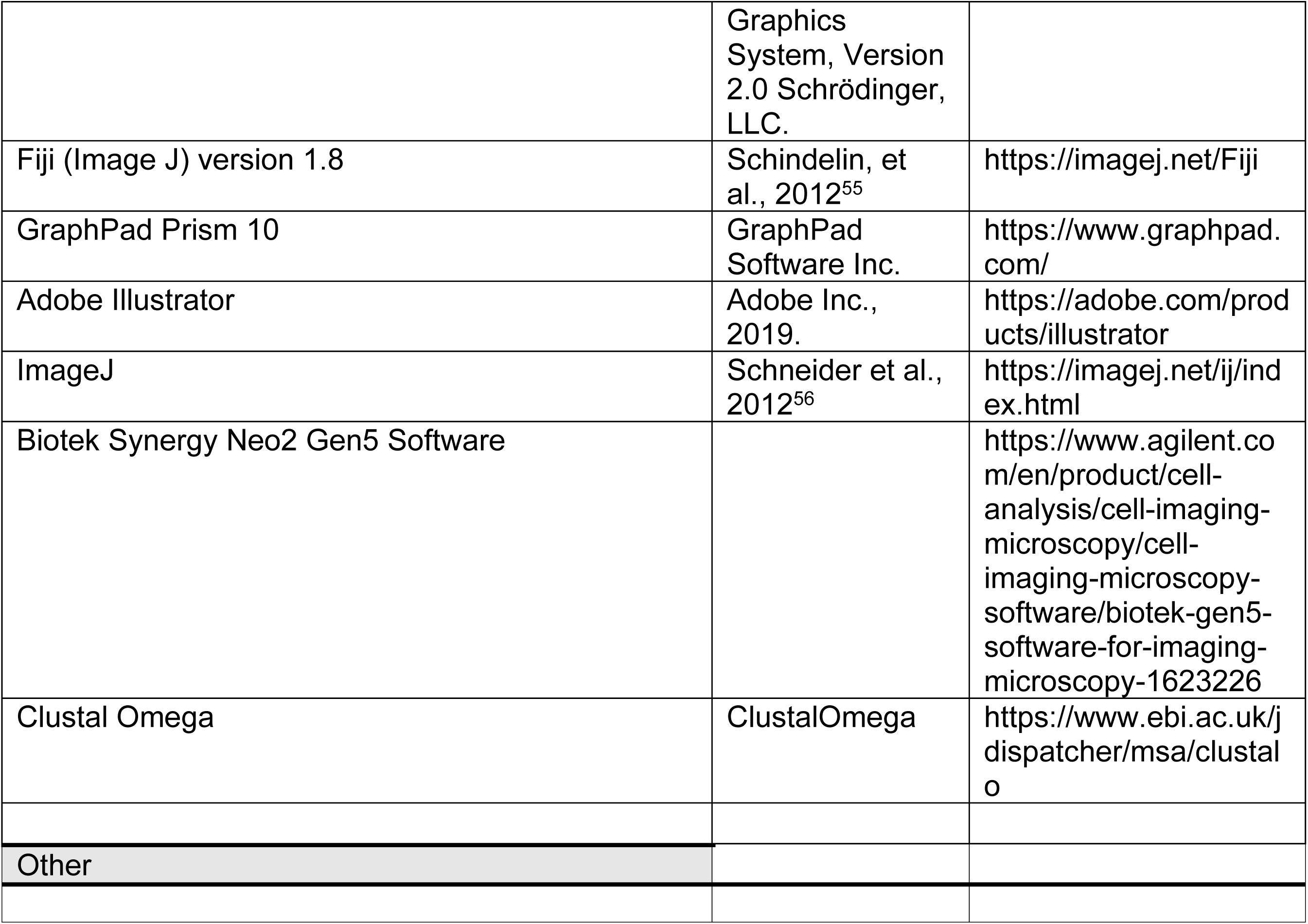

## Notes

### Competing Interest Statement

The authors have declared no competing interest.

