## Supplementary figures and images for "Inhibition of coronaviral exonuclease activity by TRIM-mediated SUMOylation"

### Supplementary Figure 1-4

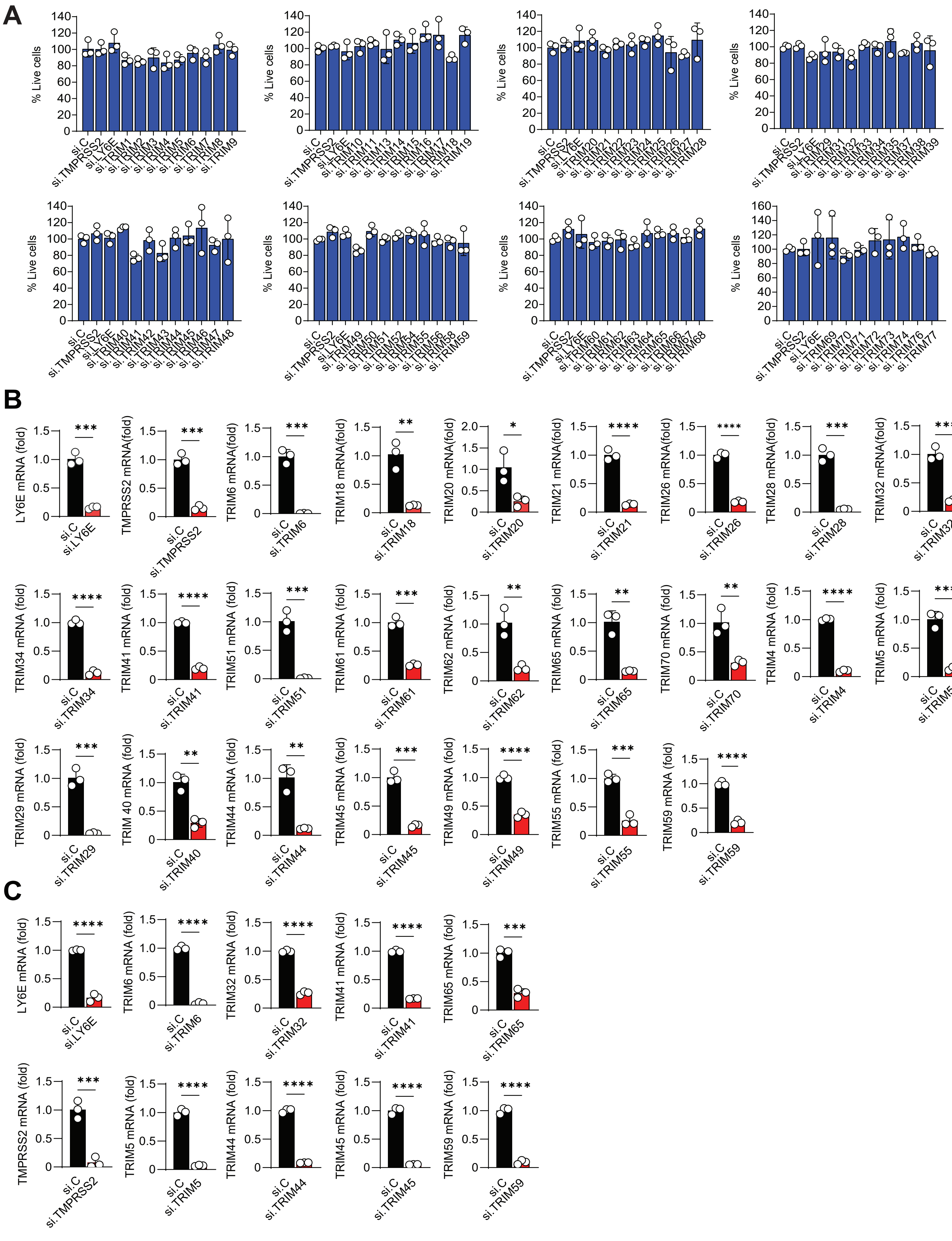

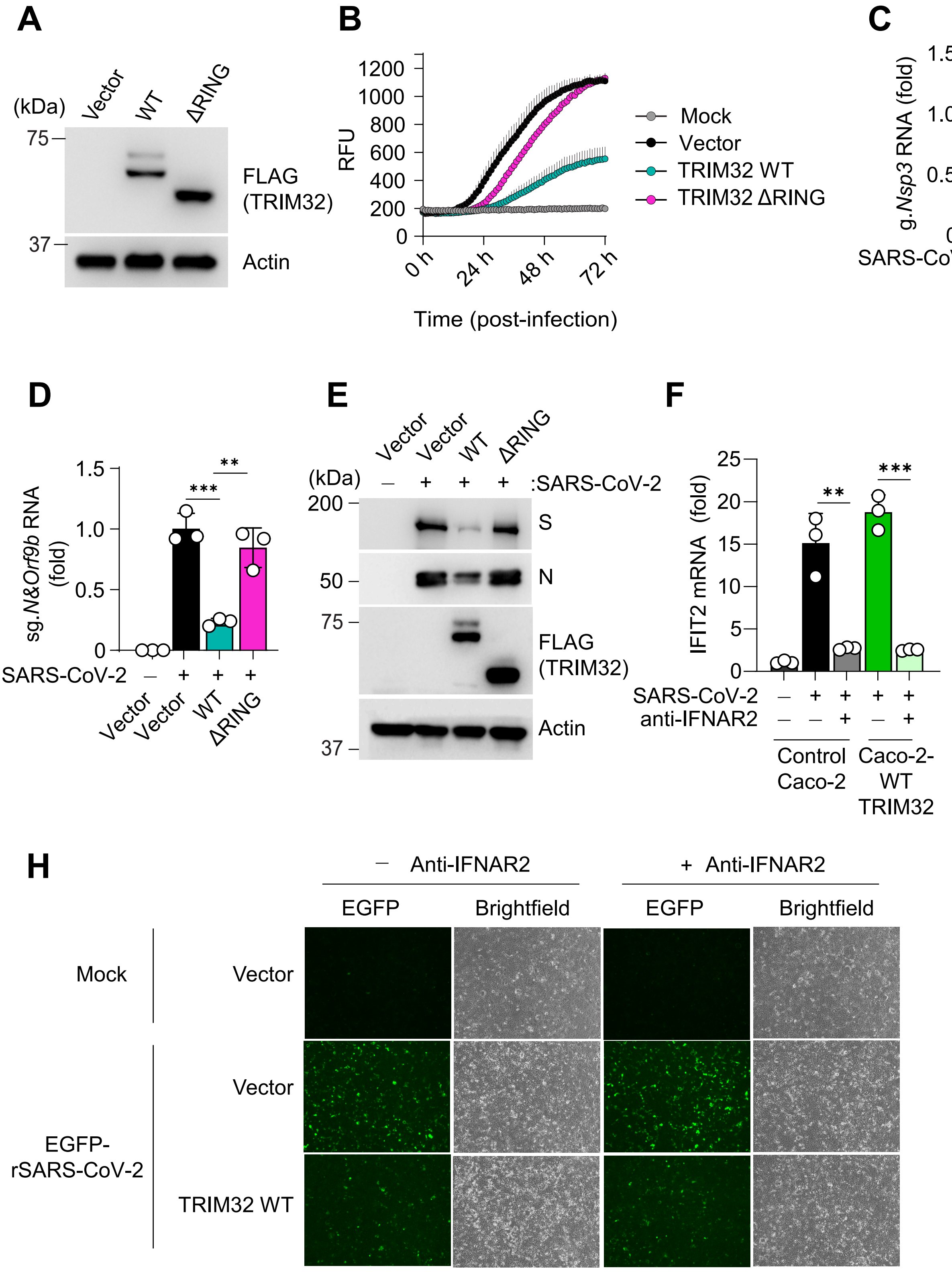

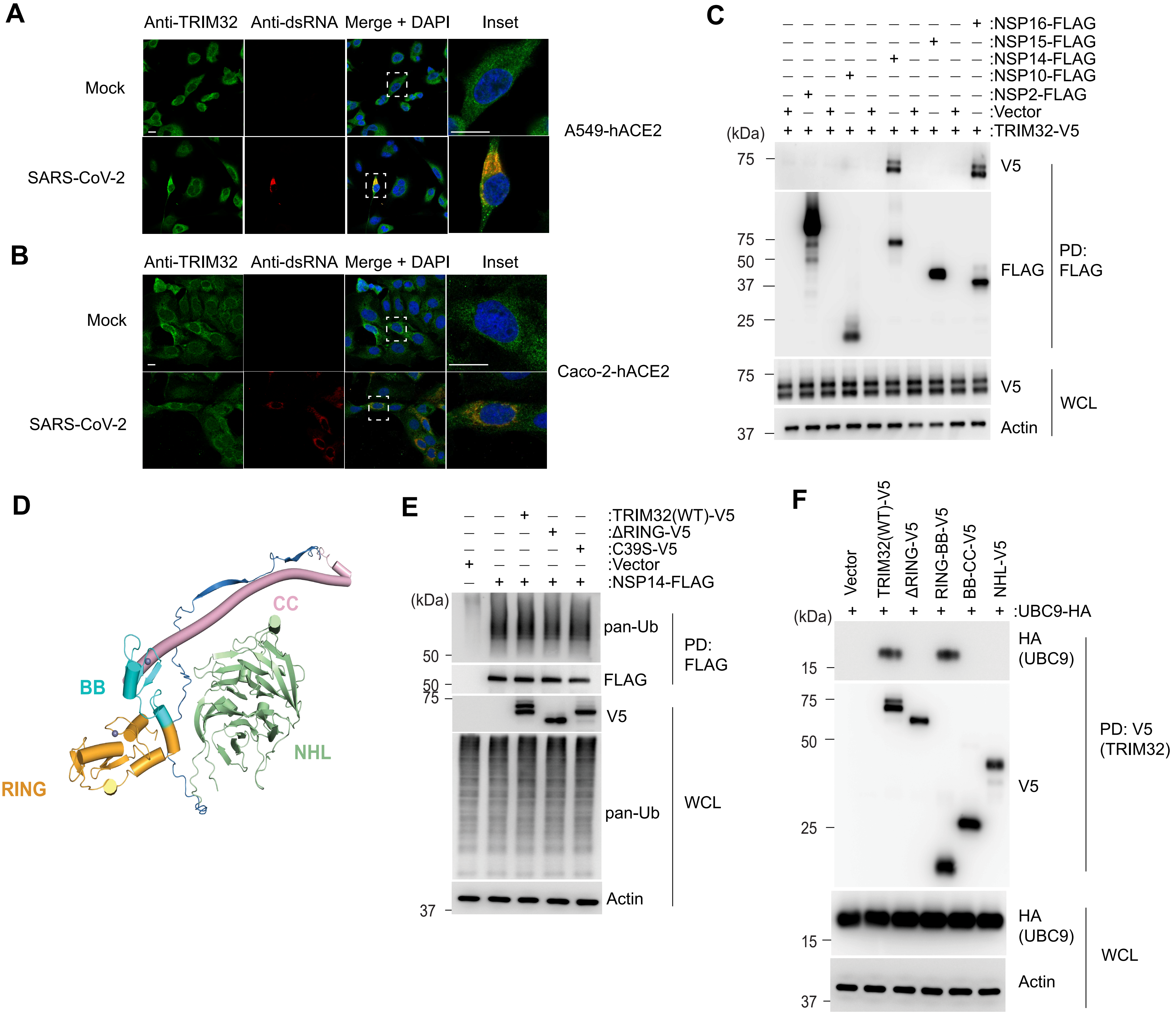

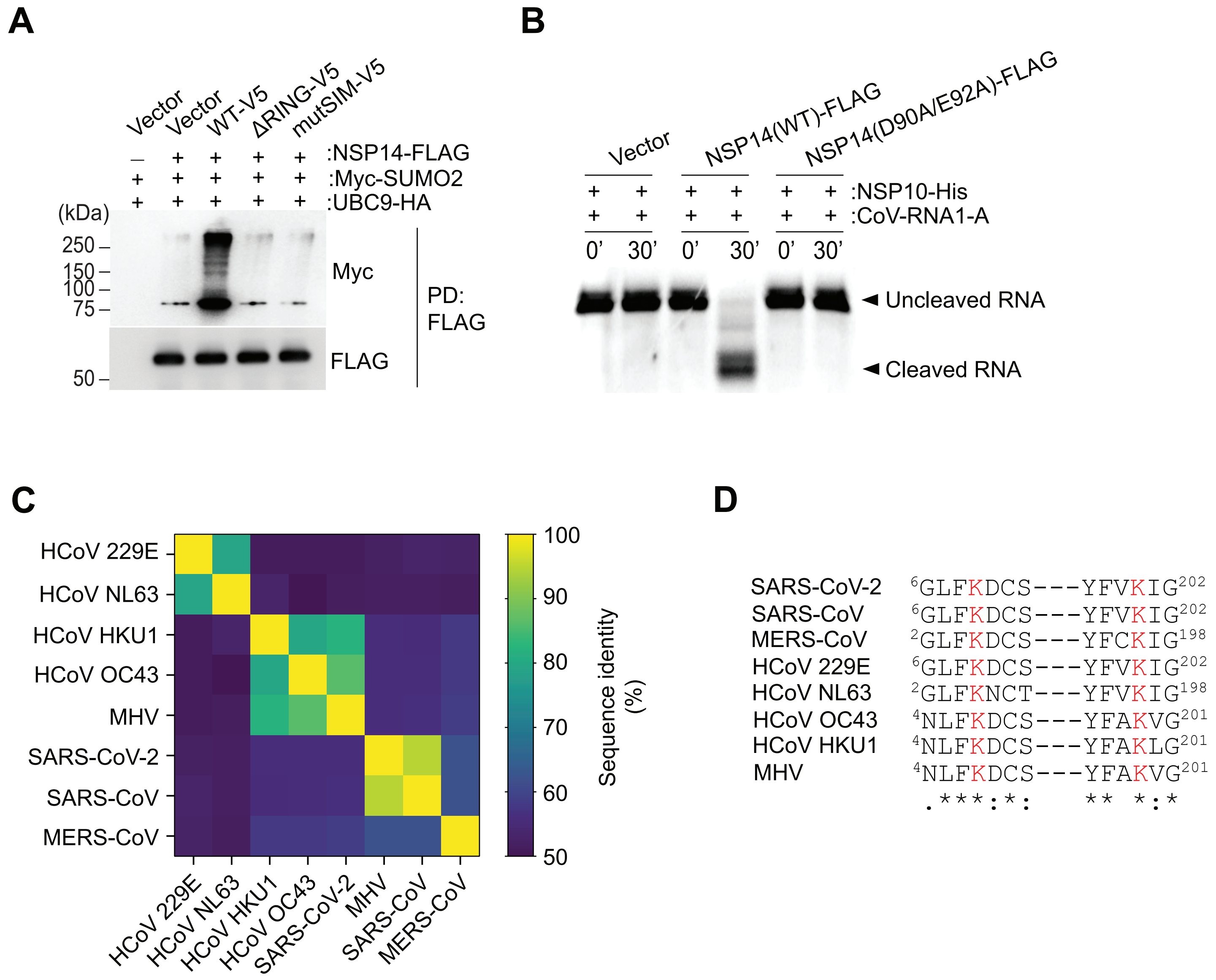
